# Variable Schwann cell merlin inactivation is targetable with TEAD1 inhibition in schwannomas

**DOI:** 10.1101/2025.11.15.688608

**Authors:** Maxwell T. Laws, Dhruval Bhatt, Debjani Mandal, Nikhil Ramavenkat, David T. Asuzu, Stefan Stoica, Ihika Rampalli, Dustin Mullaney, Liyam Laraba, Hannah Odom, Niveditha Ravindra, Sheelu Varghese, Tracy Tang, Xiyuan Zhang, John F. Shern, Abdel Elkahloun, Bayu Sisay, Dragan Maric, Kory Johnson, Zied Abdullaev, Kenneth Aldape, Ronna P. Hertzano, Hung N. Kim, David Parkinson, Prashant Chittiboina

**Author notes:** These authors contributed equally to this work (Maxwell T. Laws, Dhruval Bhatt). **Corresponding Author:** Prashant Chittiboina, MD, MPH. Tenure Track Investigator, Neurosurgery Unit for Pituitary and Inheritable Diseases, National Institute of Neurological Diseases and Stroke, National Institutes of Health. 10 Center Drive, Room 3D20, Bethesda, MD 20892-1414.

## Abstract

Schwann cell tumors occur frequently in association with the vestibular nerves, leading to sensorineural hearing loss, and brainstem compression. In humans, unilateral vestibular schwannomas (VS) occur sporadically (VS^spo^)^1^, or bilaterally with neurofibromatosis type 2 syndrome (NF2) – VS^nf2^.^2^ VS formation is driven by sub-haploid *NF2* gene dosage^3^, typically by biallelic loss.^4,5^ Loss of merlin promotes hippo/TEAD dependent transcriptional reprogramming, proliferation, and paracrine signaling that varies across time, and tumor volume.^4,6^ These variations lead to a clinically unpredictable course, and incomplete response to treatment. We hypothesized that Schwann cell merlin inactivation state determines cell-wise hippo/TEAD dependency and drives schwannoma pathogenesis. We analyzed clinical samples from VS^spo^ and VS^nf2^ with a multi-omics approach and detected variation in merlin activity within tumor Schwann cell population. We found that tumor-driving merlin-depleted Schwann cells (Schwann^mer-^) exhibited elevated hippo activity that was predominantly driven by TEAD1. In-silico TEAD1 perturbation led to a reversal to merlin intact Schwann phenotype. These findings, and tumor cell growth suppression were confirmed in NF2^fl/fl;Peri-Cre^ mouse model^7^, and in human derived schwannoma cells treated with a pan-TEAD auto palmitoylation inhibitor VT3989.^8^ Our computational and experimental results confirm that TEAD1 inhibition could be a potent, targeted strategy for schwannomas.

## Introduction

In mammalian Schwann cells, TEA domain family (TEAD) proteins interact with YAP (Yes-associated protein) and its paralogue, TAZ (Transcriptional coactivator with PDZ-binding motif) to regulate key homeostatic function including myelination.^9–11^ The TEAD/YAP-TAZ axis is regulated by merlin^12,13^, which is encoded by the *NF2* gene in humans. Schwann cell tumors (schwannomas) in humans can occur at central or peripheral nervous system nerves. Schwannomas occur frequently in association with the vestibular nerves, leading to sensorineural hearing loss, and brainstem compression. In humans, unilateral vestibular schwannomas (VS) occur sporadically (VS^spo^)^1^, or bilaterally with neurofibromatosis type 2 syndrome (NF2) – VS^nf2^.^2^ VS formation is driven by sub-haploid *NF2* gene dosage^3^, typically by biallelic loss.^4,5^ *NF2*/merlin, suppresses tumorigenesis via phosphorylation and inactivation of YAP.^14,15^ In human schwannomas, a near universal loss of merlin is typically mediated either via inactivating mutations, or 22q loss.^3–5,16^ Epigenetic regulation of merlin activity or *NF2* dosing is considered unlikely. While there is some evidence of clonal variation of *NF2* dose within schwannomas^17^, sub-clonal mechanisms likely explain the phenotypic or proliferative variability within tumors – intra-tumoral heterogeneity.^6,18^

Intra-tumoral heterogeneity often underlies the potential for recurrence, and resistance to targeted treatments in human tumors. Although, the exact mechanisms of intra-tumoral heterogeneity in VS remain unknown, heterogeneity is found at transcriptional, protein expression, and cellular proliferative potential levels.^19–21^ VS^spo^, may arise from one or two hits at the NF2 locus^4^, and cellular merlin expression varies significantly even across the same tumor. In patients with NF2, a majority of VS^nf2^ contain a single second-hit at the *NF2* locus^16,22,23^, and some VS^nf2^ may be polyclonal in origin.^24^ We hypothesized that apparent intra-tumoral heterogeneity of Schwann cell transcriptome hides strong signals of cellular selection and fitness that could have a potential in designing targeted therapies for Schwannomas.

Here, we analyzed VS^spo^ and VS^nf2^ with a multi-omics approach to understand the biological implications of intra-tumoral heterogeneity. We leveraged the wide spectrum of *NF2*/merlin status between *NF2*-intact (non-Schwann cells in VS^spo)^, - haploid (non-Schwann cells in VS^nf2^), and -lost (tumor driving Schwann cells in VS) to establish a distinct transcriptomic signature of merlin activity. We then found that Schwann cells with the lowest merlin activity drive tumor progression via paracrine, macrophage chemotactic, and angiogenic signaling. We found that transcription factor TEAD1 drives tumorigenic signaling in merlin depleted cells, and that effect that can be reversed by TEAD1 inhibition. We used an emerging TEAD1 inhibitor (VT3989)^8^ to show targeted activity against *NF2*-loss Schwann cells in mice^7^, and in human-derived VS tumor cell cultures.

## Results

### VS^nf2^ and VS^spo^ share common Schwann cell states

We performed a multi-omic analysis of VS^nf2^ (n=10; 8 patients), and VS^spo^ (n = 6) with overlapping single cell transcriptomic, chromatin accessibility, and spatial studies (**Fig. 1A**). Each sample was associated with detailed phenotypic and genotypic data (**Fig. 1B; Sup. Table S1**). Clustering and uniform manifold approximation and projection (UMAP)^25^ mapping did not reveal sample-specific distribution bias (**Fig. 1C**), allowing us to apply a robust annotation strategy for the single cell (scRNAseq; 15,500 cells) and single nucleus (snRNAseq; 77,930 cells) datasets (**Fig. 1D; Sup. Table S2**). For spatial RNA in-situ hybridization (RNAish; RNAscope, ACDBio, USA; 36,699 cells), multiplexed immunohistochemistry (mIHC; 2,721,272 cells), and high-resolution in-situ hybridization based spatial transcriptomic (stRNAish; Xenium, 10X Genomics, USA; 1,219,325 cells), we applied annotation strategies informed by snRNAseq dataset (**Sup. Figs. S1A and S1B**; **see Methods**). To understand the effect of assay preparation on downstream analysis, we performed parallel scRNAseq and snRNAseq in four tumors (**Sup. Fig. S1C**). Single nucleus assays captured the cell class membership (**Sup. Fig. S1D, Sup. Table S3**) with higher accuracy, and were enriched for ribosomal genes (**Sup. Fig. 1E**). However, clustering and cell-classification was unhindered (**Sup. Fig. 1C**), allowing us to integrate datasets for downstream analysis.

**Figure 1:**
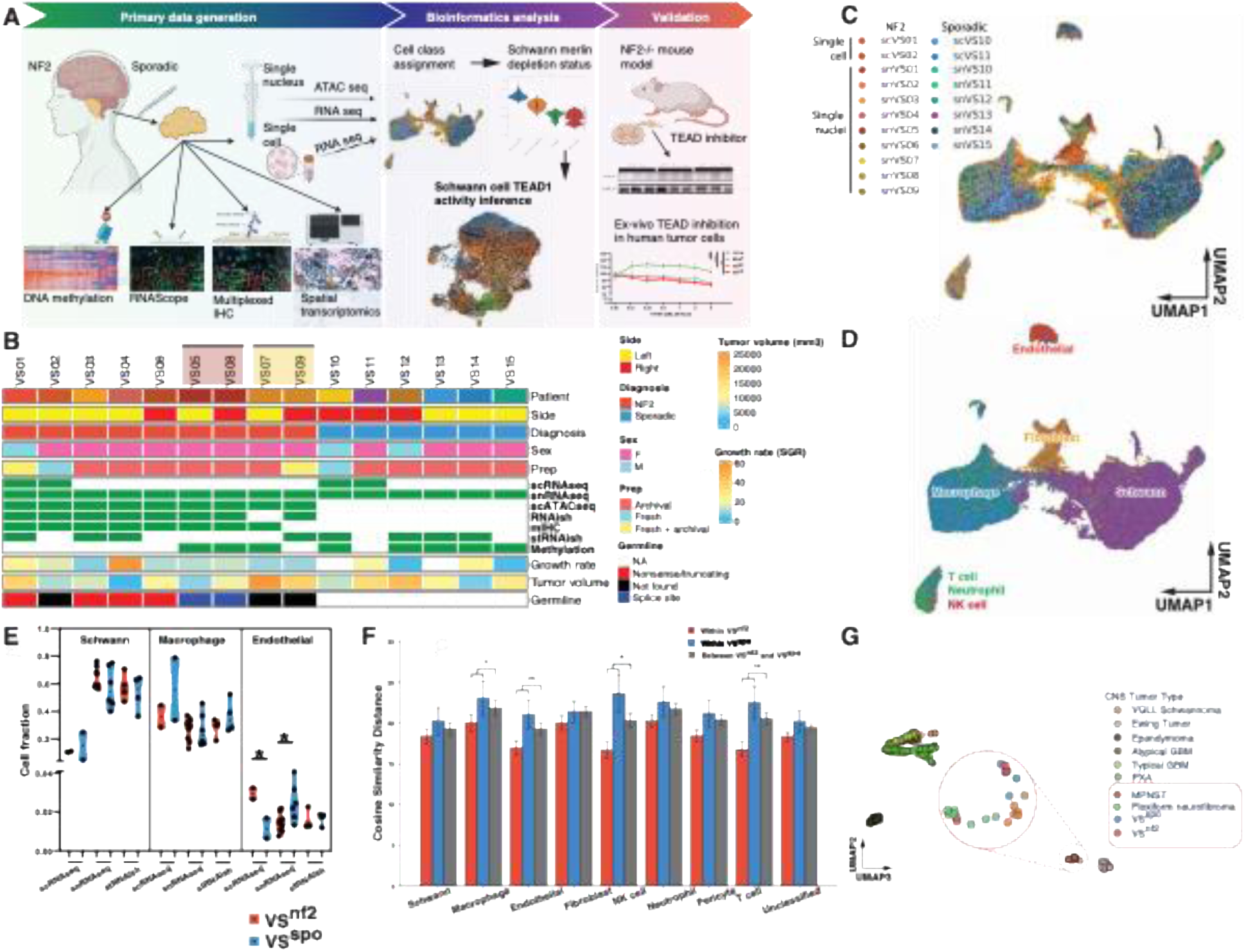
NF2 related and sporadic vestibular schwannomas share common tumorigenic pathways. **A.** Study schema. Vestibular schwannomas (NF2-related – VS^nf2^ and sporadic - VS^spo^) were surgically resected and processed for single cell and single nucleus RNAseq (sc/snRNAseq), single nucleus assay for transposase-accessible chromatin (snATACseq), multiplexed immunohistochemistry (mIHC), RNA in-situ hybridization (RNAish), chip-based bulk DNA methylation, and high-resolution spatial transcriptomic (stRNAish) analysis. **B.** Summary of human surgical samples, patient demographics, assays performed (as in A), VS tumor growth rate (specific growth rate – SGR, calculated as 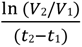 where *V* is tumor volume measured at two time (t) points.), VS tumor volume (mm^3^) and germline genotype. Two patients with NF2 contributed tumor pairs VS05-VS08, and VS07-VS09. **C.** Integrated uniform manifold approximation and projection (UMAP) with scRNAseq and snRNAseq data mapped to individual patient samples. **D.** Integrated UMAP projection with canonical cell classes (see also Sup. Table S2) **E.** Violin plot illustrating cellular fractions for VS^nf2^ and VS^spo^ with each point representing a tumor (ANOVA with Dunnet correction for multiple comparisons, p < 0.05 *) **F.** Transcriptomic cosine similarity distance of canonical cell classes between VS^nf2^ and VS^spo^ (Mann-Whitney; p < 0.05 *, p < 0.01 **). **G.** Beta values for bulk DNA methylation of CNS tumors represented on a UMAP following dimensionality reduction.

Following assay validation, we asked if VS^nf2^ and VS^spo^ could be driven by divergent mechanisms. For genotyping, we performed germline whole genome sequencing for patients with VS^nf2^ (**Sup. Table S1**). The coarse-grained UMAP distribution appeared unbiased (**Sup. Fig. S2A**), with comparable cell class memberships (**Fig. 1E**) across assays. Transcriptomically, Schwann cells from VS^nf2^ and VS^spo^ were indistinguishable (**Fig. 1F, Sup. Fig. S2B**), however, macrophages and other stromal cells demonstrated variability across the germline genotypes. Patients with NF2 are haploid for NF2/merlin activity, however Schwann cells acquire biallelic *NF2* loss in VS regardless of the germline status. With inferred copy number analysis we found a similar distribution of 22q-/-Schwann cells (**Sup Fig S2A**) in VS^nf2^/VS^spo^, however, we found discrepancies in macrophage population distribution (**Sup Figs S2C, and S2D**). VS^nf2^ were enriched for M2-like macrophages^26^ (**Sup. Figs. S2E, and S2F**), that demonstrated dysregulation of hippo, and cell-adhesion pathways (**Sup Fig S2G**), likely due to their *NF2* haploid status. Next, we analyzed whether epigenomic patterns could distinguish VS^nf2^ and VS^spo^. With bulk DNA methylation analysis, we found that VS^nf2^ and VS^spo^ were located within a subcluster of Schwann tumors (**Fig. 1G**) and clustering did not separate VS^nf2^ from VS^spo^ (**Sup. Fig. S2H**). When comparing VS^nf2^ with VS^spo^, we found that Schwann cells were indistinguishable, however the germline haploid *NF2* dosage appeared to result in an M2-like macrophage phenotype in VS^nf2^. In subsequent analyses, we focused on Schwann cell states across all VS that could be driving tumor progression.

### Merlin dosage determines Schwann cell states

Mammalian cells are sensitive to *NF2*/merlin^27^ dosage. *NF2* haploinsufficiency and/or merlin depletion deterministically affects the cellular phenotype. In schwannomas, chromosome 22q loss (via loss of heterozygosity or biallelic deletion) is not always the mode of NF2/merlin inactivation, making the identification of Schwann cells with merlin depletion challenging. To overcome this, we first defined a consistent transcriptomic signature of *NF2*/merlin loss (**Fig. 2A**) that was independent of the underlying genetic mechanism. Leveraging tumors with a distinct signal for chromosome 22q loss (e.g., snVS03 and snVS13), we identified 158 genes that were consistently upregulated in Schwann cells and used these to derive a merlin depletion score (MDS) (**Fig. 2B**). Across tumors, the MDS correlated strongly with hippo signaling pathway^28^ and *NF2*-knockout cell-line^29^ derived scores (r = 0.46–0.70) but not with 22q copy-loss status (r = –0.40), indicating that it reflects merlin functional depletion rather than chromosomal dosage (**Sup. Fig. S3A; Sup. Table S4**). The MDS showed improved discrimination for identifying Schwann cell populations with merlin loss compared to the hippo pathway activation score^28^, and transcriptomic signatures derived from experimental *NF2* knockout cells (HS01, HS11, Syn01, Syn05)^29^ (**Sup. Figs. S3B and S3C**). To evaluate how coherently each score mapped onto the single-cell manifold, we computed Moran’s I score using the UMAP embedding as a generic spatial framework.^30^ The MDS consistently displayed the highest autocorrelation across tumors (mean Moran’s I = 0.84 ± 0.06), exceeding values for hippo, HS01/HS11, and Syn01/Syn05 (**Sup. Fig. S3D**). Taken together, these results demonstrate that the MDS provides the most discriminative and biologically coherent measure of merlin functional loss across both VS^nf2^, and VS^spo^, and it was therefore used for downstream analyses.

**Figure 2:**
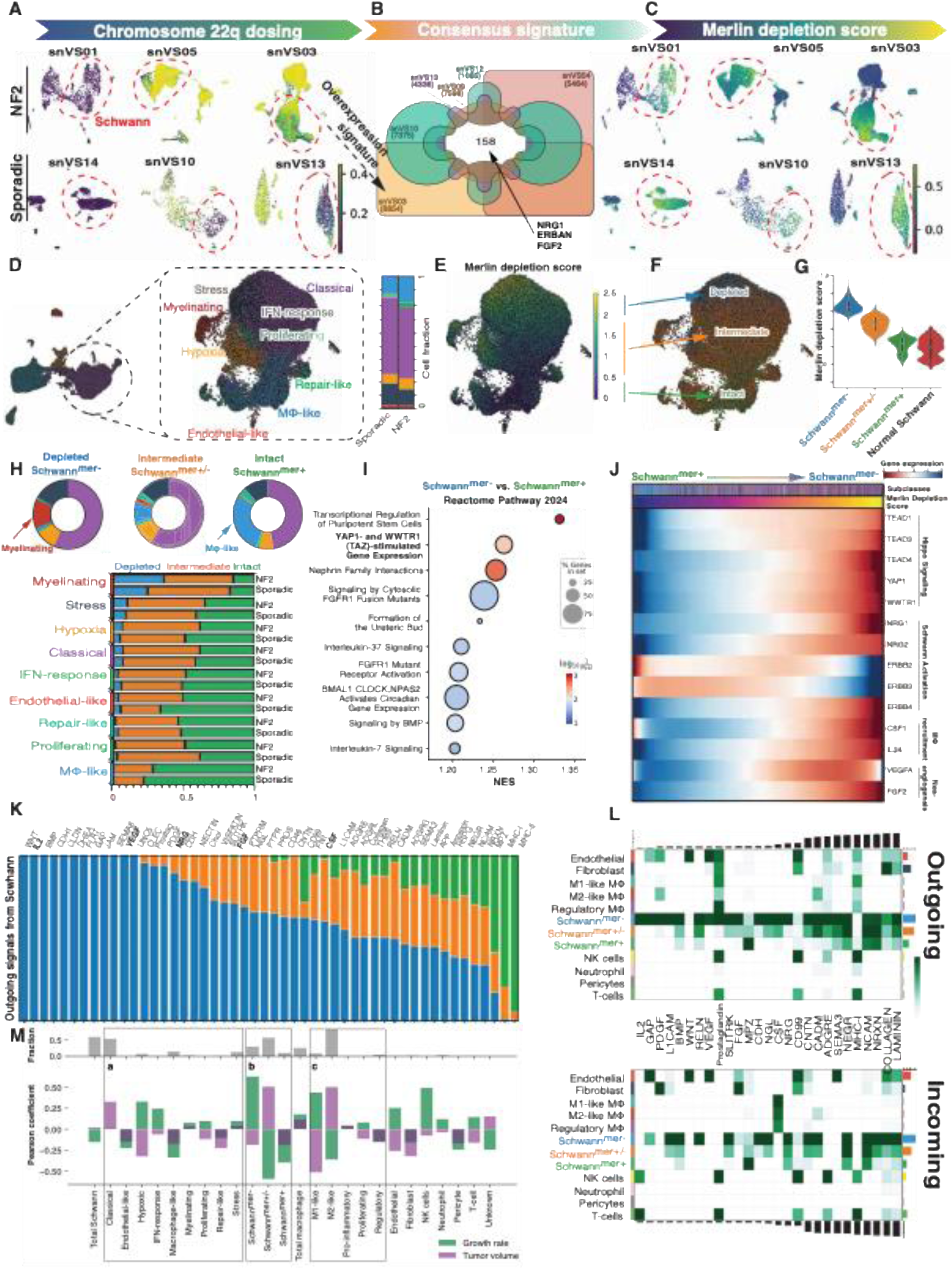
Variable Schwann cell states are detectable as merlin depletion score. **A.** Copy number estimation for 22q region mapped over uniform manifold approximation and projection (UMAP) for individual tumor samples. Representative VS^nf2^ and VS^spo^ are shown to demonstrate the range in detection of chromosome 22q loss. Tumors demonstrating distinct chromosome 22q loss are shown on the right. Schwann cell clusters are highlighted with dashed red lines. **B.** Venn diagram of differentially over-expressed genes found in Schwann cells with a distinct 22q loss signature (such as snVS03 and snVS13 in A). Merlin depletion score (MDS) generated from the overlapping sets of genes in 6 tumors with distinct 22q loss signatures (n = 158 genes). **C.** MDS applied to the same single RNAseq UMAPs generated in A. Schwann cell clusters are highlighted with dashed lines. **D.** Schwann cell sub setting, and sub-classification shown with UMAPs. The relative proportions of sub-classes are plotted in the adjacent bar-graph. Bars retain the sub-class specific colors from shown in UMAP. **E.** MDS applied to Schwann cell UMAP. **F.** Reclassification schema for Schwann cells into Schwann MDS classes using standard deviation (σ) cutoffs (depleted cells >2σ, intermediate 2σ< x >2σ, intact <2σ). **G.** Schwann^mer-^, Schwann^mer+/-^, and Schwann^mer+^ cell nomenclature applied to depleted, intermediate and intact cells respectively (from F). Schwann MDS is plotted with violin plots, and compared with MDS of normal Schwann cells. **H.** Sub-classification of Schwann cells according to transcriptomic profile. Pie charts at the top represent the relative proportions of sub-classes to Schwann MDS classes. Bar chart below represents the same dataset, with Schwann MDS class portions shown per Schwann sub-class. **I.** Top differentially expressed pathways analyzed with gene set enrichment analysis (gene set – Reactome Pathway 2024) represented as a dot plot. **J.** A heatmap for tumorigenic genes arranged along the Schwann MDS pseudotime. **K.** Major outgoing signals from Schwann cells inferred from ligand-receptor analysis. Colors represent the relative fraction of signal per Schwann MDS class per signal. **L.** Major outgoing and incoming signals inferred from ligand-receptor analysis. Rows represent the major cell classes, and columns represent the signaling pathways plotted. Row and column aggregates are plotted as marginal bar plots. **M.** Correlation analysis (Pearson correlation coefficient) of cell class, and sub-class and Schwann MDS class fractions with vestibular schwannoma growth rate, and tumor volume at surgery. Insets represent Schwann cell sub-classes (a), Schwann MDS classes (b), and macrophage subclasses (c).

The Schwann cells were then subset, re-clustered, and sub-classified according to the dominant gene expression patterns (**Sup. Fig. 4A, Sup. Tables S5**), that matched with previous Schwann cell sub-classification schema^31^ (**Sup. Fig. 4B**). There were no significant differences in sub-class membership between VS^nf2^ and VS^spo^ (p = 0.5, Mann-Whitney test) (**Fig. 2D**). We noted a gradient in the MDS across the Schwann cells (**Figure 2E**), allowing us to re-classify cells according to quartiles for MDS: +σ > 2σ > -σ as merlin-depleted (Schwann^mer-^), merlin-intermediate (Schwann^mer+/-^), and merlin-intact (Schwann^mer+^) (**Fig. 2F**). The transcriptomes of publicly available normal Schwann cells^18,32,33^ (2,395 cells) were confirmed to have a Schwann^mer+^ MDS signature (**Fig. 2G**).

The myelinating subset of Schwann cells were the largest, and macrophage-like (MΦ-like) cells were the smallest sub-class of Schwann^mer-^(**Figure 2H**). Hippo signaling (via YAP/TAZ stimulated gene expression)^14,15^ pathway was highly upregulated in Schwann^mer-^ (**Fig. 2I, Sup. Table. S6**). We found highly coordinated elevations in hippo signaling (*TEAD*s, *YAP1*, *WWTR1*), Schwann cell activation (*NRG1*, *NRG2*), macrophage recruitment (*CSF1*, *IL34*), and angiogenesis (*VEGFA*, *FGF2*) along the MDS pseudotime (**Fig. 2J**), suggesting a congruence between MDS and tumorigenic signaling driving Schwannoma progression.^34–36^ Myelinating Schwann sub-class appeared to drive major outgoing signaling, while the hypoxia Schwann sub-class was driving VEGFA signaling (**Sup. Fig. 4C).** Ligand-receptor interaction analysis revealed that Schwann^mer-^ contributed to a majority of the outgoing signals from Schwann cells to the VS microenvironment (including IL2, VEGF, NRG, and CSF1 signaling) (**Fig. 2K**). Schwann^mer-^ were promiscuous senders and receivers, with other receivers being Schwann^mer+/-^, Schwann^mer+^, endothelial cells and macrophages (Mφ) (**Fig. 2L**). Clinically, Schwann^mer-^ abundance in VS was highly correlated with tumor growth rate (Pearson coefficient = 0.61) (**Fig. 2M**). Taken together, our findings suggest that merlin depletion in a subset of Schwann cells could be a primary driver of VS tumor progression.

### Nuclear retention of NF2 mRNA is related to high MDS in Schwann cells

Merlin immunoreactivity is a poor indicator of the underlying mechanisms of *NF2*/merlin inactivation.^37^ To overcome the limitations of protein-level^31,38^, and spot-aggregated spatial transcriptomics^39^, we investigated VS tissues with high-resolution spatial transcriptomics platform (stRNAish). Using a custom 300-gene probe-set (**Sup. Table S7**), we investigated the sub-cellular localization, and consequences of NF2/merlin inactivation spatially. Following label-transfers from snRNAseq datasets^40^ (**Sup. Fig. S5A**), we identified the canonical cell classes, Schwann/macrophage subclasses, and Schwann merlin status (VS^nf2^, n=4; VS^spor^, n=4) (**Fig. 3A**). We had noted that neither chromatin accessibility at the *NF2* locus (**Sup. Fig. S5B**) nor *NF2* transcript (**Sup. Fig. S5C**) expression was associated with Schwann MDS at the single cell level. We asked if nuclear retention of the *NF2* transcript^41^ played a role in merlin gene expression status and MDS. We found that Schwann^mer-^ retained nuclear NF2 mRNA compared to Schwann cells with lower MDS (**Fig. 3B**). Tumor cells frequently demonstrate enlargement of nuclear size^42^, a phenomenon we observed in increasing nuclear size in Schwann cells with increasing MDS (**Fig. 3C**), but, without a change in total cell size. We found that global chromatin discompaction of Schwann^mer-^ compared to Schwann^mer+^ cells in snATACseq dataset (**Sup. Fig. 5D**); a phenomenon that could underlie nuclear expansion. Correspondingly, we found that the nuclear retained mRNA in Schwann^mer-^ was significantly enriched for RNA preprocessing (**Sup. Fig. S5E**).

**Figure 3:**
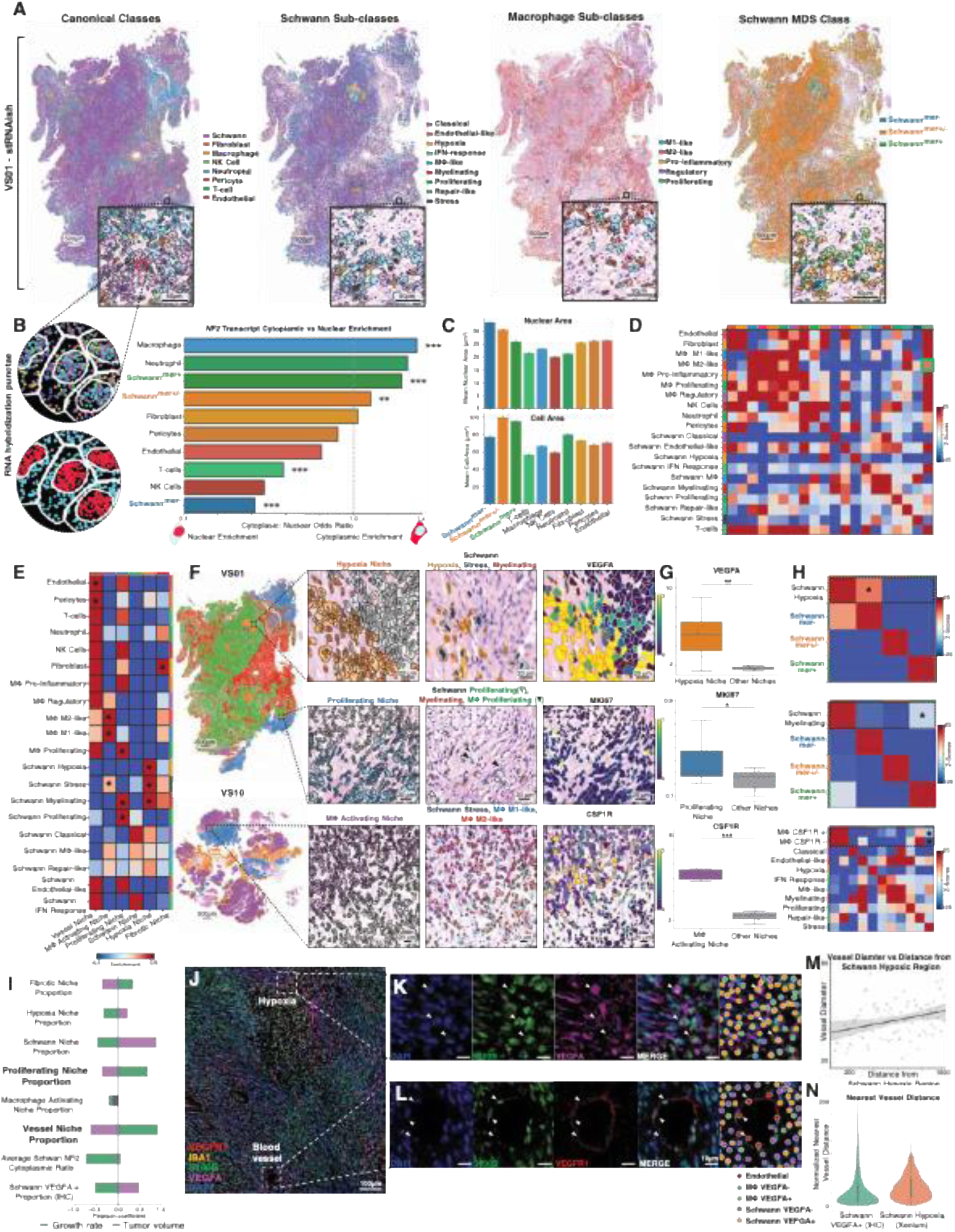
Schwann MDS classes occupy spatial niches to drive Schwannoma tumor progression. **A.** Representative high-resolution spatial transcriptomic (stRNAish) map of a tumor (VS01) with canonical cell classes, Schwann sub-classes, macrophage sub-classes, and Schwann MDS classes overlaid (scale bar = 500µm). Insets in each panel are magnified view demonstrating the cells outlined, and mapped over a hematoxylin & eosin background (scale bar = 50µm). **B.** Subcellular compartmentalization of *NF2* transcripts. Left: cell schematic showing all transcripts (top) and classification of nuclear (red) vs cytoplasmic (blue) transcripts (bottom). Center: relative nuclear-to-cytoplasmic enrichment ratio for *NF2* transcripts across Schwann MDS classes and canonical cell types (values < 1 indicate greater nuclear enrichment; > 1 indicate greater cytoplasmic enrichment relative to other cell types). Asterisks denote significance (p < 0.05 *, p < 0.01 **, p < 0.001 ***). Right: bar plots (mean ± SEM) of nuclear area (top) and cell area (bottom). **C.** Bar plots summarizing the nuclear area (top), and total cell area (bottom) of Schwann MDS classes, and stromal cells in the stRNAish data. **D.** A clustered heatmap of spatial neighborhood enrichment (z-scores) for Schwann sub-classes and stromal cells in the integrated dataset generated from stRNAish data. The green dashed squares highlight high enrichment probabilities for M2-like macrophages with repair-like Schwann cell sub-class, and of T cells with interferon response Schwann sub-class. **E.** Heatmap plot for enrichment of cell subtypes within spatial niches (Schwann, hypoxia, proliferating, macrophage-activating, fibrotic, vessel); asterisks (*) indicate key cell subclasses within each niche. **F.** Representative spatial niche overlays on stRNAish datasets (VS01, a VS^nf2^, top; VS10, a VS^spo^, bottom). Color code for spatial niches is the same as niche title hues and Sup. Fig. 5G (scale bar = 500µm). Inset panels demonstrate the spatial niches (left), Schwann sub-classes (middle), and gene expression of key niche-specific genes (right). Color bars on the right indicate the expression levels in gene expression panels (scale bar = 20µm). **G.** Box and whisker plots summarizing niche-specific gene expression: *VEGFA* (hypoxia vs other niches, top), *MKI67* (proliferating vs other, middle), and *CSF1R* (macrophage-activating vs other, bottom). Each point represents an averaged niche-specific expression per tumor. Asterisks denote significance (p < 0.05 *, p < 0.01 **, p < 0.001 ***). **H.** Heatmaps of neighborhood-enrichment of hypoxia Schwann sub-class relative to Schwann MDS classes (top), myelinating Schwann sub-class relative to Schwann MDS classes (middle), and CSF1R-positive vs CSF1R-negative macrophages relative to Schwann sub-classes; asterisks (*) mark key interactions reported in the main text. I. Bar plot summary (Pearson coefficient) of clinical tumor characteristics and their relationships with the stRNAish spatial variables (e.g., niche proportions, average Schwann NF2 transcript cytoplasmic ratio, Schwann VEGFA positivity) in each VS tumor. **J.** Representative merged multiplexed immunohistochemistry image from VS01 tumor. Primary antibody probe sub-titles in the respective hues. Hypoxia and vessel niches are marked by squares (scale bar = 100µm). **K and L.** Inset panels representing hypoxia spatial region (top), and vessel spatial region. Subtitles indicate the probes in respective hues. White arrowheads mark Schwann VEGFA+ cells (top), and endothelial cells (bottom) (scale bar = 10µm). **M.** Scatterplot mapping the blood vessels diameter (y-axis) in µm, and their distance from the centroids of hypoxia spatial niche (from stRNAish datasets) in µm. Each point represents a single blood vessel. Correlation and 95% confidence interval is overlaid on points (Pearson’s correlation, r = 0.30, p < 0.01). **N.** Violin plots summarizing the relationship of within-sample normalized distance of blood vessels from Schwann VEGFA+ cells (mIHC), and hypoxia Schwann sub-class cells (stRNAish).

**Figure 4:**
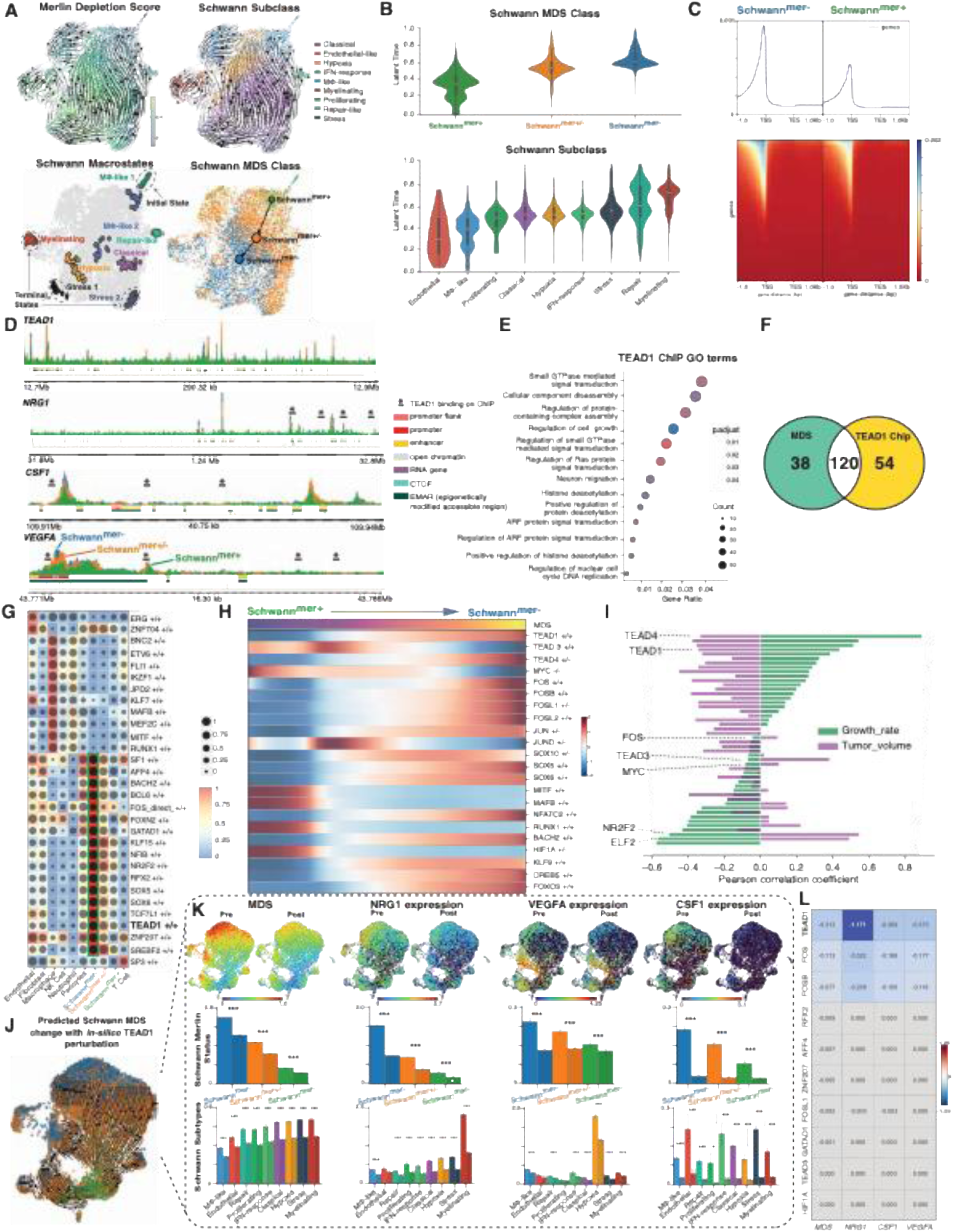
Schwann merlin depletion occurs as a plastic transcriptional cell state dependent on TEAD1 activity. **A.** Representative uniform manifold approximation and projection (UMAP) of Schwann cells from a single tumor sample (VS04), overlaid with RNA velocity trajectories. Top left: RNA velocity vectors colored by merlin depletion score (MDS). Top right: UMAP colored by Schwann sub-classes. Bottom left: inferred Schwann sub-class macrostates, indicating initial and terminal states based on RNA velocity. Bottom right: Directional PAGA graph abstraction of RNA velocity across Schwann cells stratified by MDS status. **B.** Violin plots of Schwann MDS classes (top) and Schwann sub-classes (bottom), ordered by increasing latent time derived from RNA velocity–based pseudotime analysis. **C.** Heatmap plot depicting chromatin accessibility at genes between Schwann^Mer−^ and Schwann^Mer+^ cells. Chromatin regions are mapped from 1 kilobase (kb) upstream of transcription start site (TSS) through 1 kb downstream of transcription end site (TES). **D.** Single nucleus assay for transposase accessible chromatin (snATACseq) coverage plots for selected genes across Schwann MDS classes. **E.** Gene Ontology (GO) GSEA analysis of TEAD1 chromatin immunoprecipitation (ChIP) targets in Schwannoma samples (n = 3). **F.** Venn diagram summarizing the overlap between genes contributing to the MDS, and TEAD1 ChIP targets summarized in E. **G.** Heatmap displaying inferred transcription factor (TF) activity across Schwann cells, stratified by Schwann MDS classes and canonical cell types. Color in each cell represents average target gene enrichment, and dot size indicates average target region accessibility. **H.** Smoothed heatmap of selected inferred TF activity trends across the Schwann MDS class pseudotime. **I.** Bar plot showing correlations (Pearson coefficient) between selected average TF activities and clinical metrics (tumor growth rate and volume). **J.** UMAP visualization of Schwann cell Merlin status with arrows showing predicted effects of TEAD1 transcription factor activity perturbation on Schwann MDS class phenotype change (*in-silico* knockout, lowest expression set to zero, five simulation iterations). **K.** Comparative visualization of pre- and post-TEAD1 perturbation (as in J.) states showing MDS and expression changes in selected genes across Schwann MDS and sub-classes (Wilcoxon rank sum test; (p < 0.05 *, p < 0.01 **, p < 0.001 ***)). **L.** Heatmap showing predicted log2FC in MDS, *NRG1*, *CSF1*, and *VEGFA* with *in-silico* perturbation of top active transcription factors. Each row represents a transcription factor tested.

**Figure 5:**
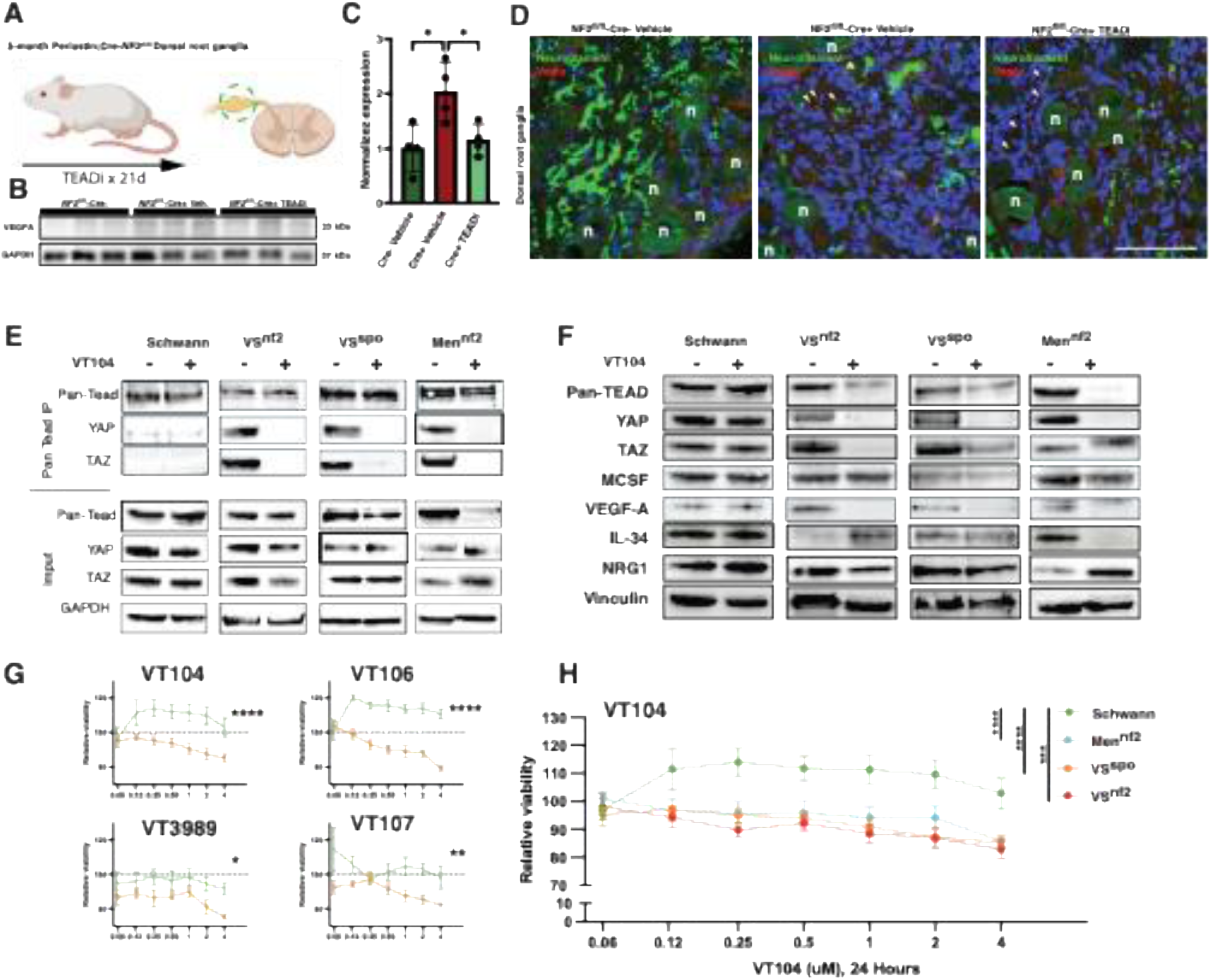
Pharmacological TEAD1 inhibition reverses merlin depletion mediated tumorigenic pathways. **A.** Summary of Periostin;Cre-NF2fl/fl mouse treatment with TEAD auto palmitoylation inhibitor (TEADi, VT3989), and dorsal root ganglia (DRGs) harvesting. **B.** Representative western blot of 5-month DRGs for VEGFa expression in Cre-littermate control mice (NF2fl/fl-Cre-; lanes 1-3), NF2fl/fl-Cre+ vehicle treated mice (NF2fl/fl-Cre+ Veh; lanes 4-6) and NF2fl/fl-Cre+ mice treated with 30 mg/kg of TEADi for 21d (NF2fl/fl-Cre+ TEADi; lanes 7-9). Blot shows 3/4 biological repeats, GAPDH was the loading control. **C.** Quantification of data represented in A, presented as mean ± SEM, one-way ANOVA with Bonferroni’s multiple comparisons test. P < 0.05 (*). **D.** Representative immunofluorescence images of DRGs demonstrating the distribution of neurofilament (green) marked neuronal cell bodies (n), and vegfa (red) within Schwann cells marked with white arrowheads (scale bar = 100 µm). **E.** Co-immunoprecipitation of human normal Schwann cells, and primary cells cultures from NF2 related vestibular schwannoma (NF2 Swn), sporadic vestibular schwannoma (Spo Swn), and NF2 related intracranial meningioma (NF2 Men). The top section demonstrates the results from pan-TEAD immunoprecipitation with and without treatment with TEADi (VT104; 4 µM, 6 hours) for each cell type. The bottom section demonstrates the blots from cell lysates (input), with GAPDH as loading control. **F.** Representative western blot of the 4 cell types as in E., treated with TEADi – VT104 (4 µM, 6 hours), and the effects on tumorigenic mediators (MCSF, VEGF-A, IL-34, and NRG1). Vinculin was used as loading control. **G.** Comparative survival (CellTiter-Glo) of normal human Schwann cells (green), and schwannoma cells (Spo Swn; orange) in primary culture, with exposure to 4 TEADis (VT104, VT106, VT3989, and VT107), at 24 hours, across concentrations (0.06 – 4 µM). Data presented as mean ± SEM, with the results of one-way AVONA and Bonferroni correction for multiple testing. P = 0.015 (*), 0.0013 (**), < 0.0001 (****). **H.** Comparative survival of 4 human cell types as in E., following treatment with VT104 for 24 hours across concentrations (0.06 – 4 µM). Data presented as mean ± SEM, with the results of one-way AVONA and Bonferroni correction for multiple testing. P = 0.0005 (***), < 0.0001 (****).

### VS tumor progression is driven by distinct transcriptomic and protein expression spatial niches

We next explored the inter-cellular interactions between Schwann subclasses and stromal (non-Schwann) cells that could underlie progression of VS. Cell-level neighborhood enrichment analysis^30^ revealed that M2-like macrophages were closely associated with repair-like Schwann subclass, while T-cells were closely associated with IFN response Schwann subclass (**Fig. 3D**). We expanded the analysis and identified 6, non-overlapping spatial niches^43^ across VS specimens that demonstrated relative enrichment of Schwann subclasses, and stromal cells (**Fig. 3E**). We explored the hypoxic, proliferating, and macrophage activating niches further (**Fig. 3F**). In the hypoxia niche, we found enrichment of hypoxia, myelinating and stress sub-classes of Schwann cells (**Fig. 3F, top row**); sub-classes associated with high MDS scores (**see Fig. 2H**). In the hypoxia niche, we found an enrichment of Schwann cells expressing *VEGFA*, likely driving angiogenesis (**Fig 3G, top**). The proliferating niche was enriched for myelinating and proliferating Schwann sub-classes, and proliferating macrophage cells; with a corresponding enrichment of proliferating Schwann cells with high *MKI67* expression (**Fig. 3G, middle**). The macrophage activating niche was enriched with M1-and M2-like macrophages and Schwann stress sub-class; with an expected higher expression of *CSF1R* in the macrophages in this niche (**Fig 3G, bottom**).

We extended the niche analysis to explore the role of MDS in VS tumor progression. We asked which MDS classes were in proximity to Schwann sub-classes, to find that Schwann hypoxia sub-class were closely associated with Schwann^mer-^ (**Fig. 3H; top**). Meanwhile, Schwann myelinating sub-class were most closely associated with Schwann^mer+^ (**Fig. 3H; middle**), suggesting that the MDS scores are correlated with spatial niches for hypoxia, and proliferation respectively. For the macrophage proliferation niche, Schwann stress sub-class cells (expressing *CSF-1*) were in close proximity to *CSF-1R* expressing macrophages (**Fig. 3H; bottom**). Clinically, VS tumor growth-rates were correlated with the vessel niche, and expectedly, the proliferating niche. The VS tumor volume at clinical presentation was correlated with the stable Schwann cell niche (**Fig. 3I**).

We explored the relationship of the transcriptomic niches with the protein expression level landscape with multiplexed immunohistochemistry (mIHC, **Fig. 3J**). We approached this question by focusing on how hypoxic stress may lead to angiogenesis in VS tumors. Following cell classification of mIHC datasets (**see Methods**), we identified Schwann cells with VEGFA protein expression suggesting local hypoxic stress (**Fig. 3K**). The tumor blood vessels were then marked (**Fig. 3L**), and their distance from the centroids of hypoxic regions analyzed in stRNAish (**Fig. 3M**) and mIHC datasets. We found that smaller, presumably newly sprouted, blood vessels were closer to the hypoxic regions. The hypoxic regions were further removed from larger, more mature blood vessels (**Fig. 3M**), suggesting alleviation of local hypoxic pressure. Lastly, we found evidence of temporal/spatial progression in niches. When comparing the association of blood vessels with VEGFA+ Schwann cells, we found that Schwann cells with VEGFA protein expression were likely to be closer to blood vessels that the Schwann cells with *VEGFA* mRNA transcript **(Fig. 3N).** Taken together, our findings suggest that distinct vascular and proliferative spatial niches that are related to Schwann MDS mediated VS tumor growth.

### Schwann MDS is a plastic transcriptional cell state dependent on TEAD1 activity

We sought to understand if Schwann MDS was indicative of plastic transcriptional states. With a robust analysis strategy using RNA velocity^44^, and lineage tracing analysis (**Fig. 4A**)^45^, we identified macrostates across Schwann cell MDS and sub-class states. Unsupervised initial cell states were identified as Mφ-like/Schwann^mer+^ cells. Terminal states were Schwann^mer-^ cells belonging to the myelinating and stress subclasses. Broadly, lineage inference suggested directional cell state transitions between Schwann^mer+^ and Schwann^mer-^ cell states via Schwann^mer+/-^ state. Schwann MDS was aligned with the RNA velocity latent time, and consequently, the expression of hippo genes, Schwann/macrophage activating genes, and angiogenic genes (**Fig. 4B, Sup. Fig. 6**).

We extended cellular plasticity analysis by leveraging concurrent single nucleus ATACseq (snATACseq) data. We found evidence of elevated chromatin accessibility globally (**Sup. Fig. 5D**), and at genes (**Fig. 4C**) in Schwann^mer-^ cells. We investigated whether Schwann^mer-^ cells exhibited a systematic chromatin accessibility at tumor driving loci, and found elevated accessibility at promoters and enhancers for key genes (*NRG1*, *CSF1*, and *VEGFA*) and *TEAD1* (**Fig. 4D**). Since, *TEAD1* is critical for Schwann cell development^10,46^, and underpins hippo activation in tumors^15,47^, we asked if Schwann MDS was driving tumor formation via TEAD-hippo axis. We performed TEAD1 chromatin immunoprecipitation DNA sequencing (ChIP-seq) in human VS tumor primary cell cultures. We found TEAD1 binding to promoters of genes including *MYC*, *CSF1* and *VEGFA* (**Sup. Table S8**), and an overall pro-tumorigenic signaling (e.g. ‘Regulation of cell growth’ pathway) (**Fig. 4E**). A majority of the gene loci (n = 120) identified in ChIP overlapped with the genes comprising the MDS score, confirming the convergence of MDS mediated signaling on TEAD1 transcription factor (**Fig. 4F; Sup. Table S9**).

We leveraged concurrent snRNAseq and snATACseq dataset to infer transcription factor activity across the canonical cell classes and cell states in VS.^48^ We found that TEAD1 was a key transcription factor driving Schwann^mer-^ phenotype (**Fig. 4G**). Along the MDS pseudotime, inferred TEAD1 activity was highly correlated with a transition from Schwann^mer+^ to Schwann^mer-^ cell state (**Fig. 4H**). VS tumors with elevated inferred TEAD1 activity were highly correlated (Pearson correlation coefficient = 0.44) with increased growth rates (**Fig. 4I**). With *in-silico* TEAD1 activity perturbation, we found a reversal of plastic Schwann^mer-^ state towards a Schwann^mer+^ state (**Fig. 4J**). *In-silico* TEAD1 perturbation predicted a reduction in Schwann MDS across all sub-classes, and significant reductions in expression of tumor-driving genes - *NRG1*, *CSF1*, and *VEGFA* across all Schwann sub-classes (**Fig. 4K**). The effect of *in-silico* perturbation on gene expression was limited to TEAD1 (**Fig. 4L**). Taken together our findings suggest that Schwann MDS is a transcriptional cell state that is dependent on TEAD1 activity, and that suppressing TEAD1 activity could lead to a suppression of tumorigenic pathways in VS.

### TEAD1 inhibition reverses merlin-loss mediated tumorigenic pathways

Merlin loss drives tumor formation via YAP/TAZ-TEAD complex^49^, an event that is disrupted by TEAD auto-palmitoylation inhibition^50,51^. We leveraged the demonstrated anti-tumor effect of small TEAD palmitoylation inhibitors (TEADi; Vivace Therapeutics, San Mateo) on Periostin;Cre-NF2^fl/fl^ mice^7,52^. We treated 5-month-old Periostin;Cre-NF2^fl/fl^ mice with intraperitoneal TEADi for 21 days (**Figure 5A**), and harvested dorsal root ganglia as previously reported^7^. We measured the abundance of vegfa in dorsal root ganglia (**Figures 5B and 5C**). We found a two-fold increase in normalized expression of vegfa in Cre+ mice, that was reversed with TEADi treatment. We found a decrease in Schwann cell associated vegfa signal with TEADi treatment (**Figure 5D**) in dorsal root ganglia of treated mice. We did not find a change in the abundance or vegfa signal associated with the neuron cell bodies in dorsal root ganglia. These results suggested that TEADi treatment can target a key tumorigenic signal in Schwannomas with inhibition of vegfa.

We then asked if pharmacologic TEADi strategy could be translated to human patients with schwannomas and NF2-related tumors. We tested four TEADi – VT104, VT106, VT3989, and VT107 (4 µM each, for 6 hours) in a single-blind *in-vitro* experimental design. With co-immunoprecipitation, we found minimal interaction of YAP and TAZ with TEAD (anti-panTEAD pulldown), whereas, a robust interaction was seen human tumor cells in primary culture (VS^nf2^, VS^spo^, and NF2-related meningioma – Men^nf2^). TEADi effectively inhibited YAP/TAZ-TEAD interaction in human tumor primary tumor cultures (**Figure 5E**). In the same set of primary cells, we found that TEADi treatment inhibited the expression of Schwann proliferation (*NRG1*), macrophage chemotactic (*MCSF* and *IL-34*), and angiogenic (*VEGFA*) factors (**Figure 5F**). Additionally, we treated the available VS^spo^ cells cultures with VT3989 and found similar effects on YAP/TAZ-TEAD interaction, and downstream tumorigenic signal mediators (**Sup. Figs. S7A and S7B**).

In single-blind experiments, we next tested the differential efficacy of the four TEADi molecules on differential tumor cell survival *in vitro*. VT104 and VT106 demonstrated a robust therapeutic window, with minimal effects on normal Schwann cells when compared with schwannoma cells (**Figure 5G**). Both VT104 and VT106 appeared to be non-toxic to normal Schwann cells, while reducing the viability of tumor cells by 15 to 21% within 24 hours. Due to the scarcity of primary human tumor cell cultures, we then tested VT104 (4µM) on the 4 available primary tumor cell lines and found a reduction in tumor cell survival by at least 15% within 24 hours (**Figure 5H**). We found similar effects on survival of normal Schwann cells, as well as on VS^spo^ primary cell cultures with the other TEADi (VT016, VT3989, and VT107) (**Sup. Figs. S7C and S7D**). Taken together, these findings suggest that a broad class of TEAD auto-palmitoylation inhibitors may be used as targeted therapy for schwannomas, and other *NF2*/merlin loss driven human tumors.

## Discussion

We report that intra-tumoral heterogeneity of Schwann cells driven by differences in merlin activity in Schwann cells. We found that the merlin depletion score (MDS) is a transcriptomic surrogate for Schwann cell merlin activity. The MDS allowed us to identify Schwann^mer-^ cells that demonstrated the potential for driving tumor progression by at least three different mechanisms: a. paracrine Schwann cell activation of Schwann^mer+/-^ and Schwann^mer+^ cells (via *NRG1*), b. macrophage chemotaxis and activation (via *CSF1* and *IL34*)^53,54^, and c. angiogenesis (via *VEGFA*).^55–57^ MDS-based Schwann cell heterogeneity was detectable as spatial niches in both transcriptomic and protein-expression level analyses. Although, we did find evidence of polyclonality in VS^nf2^ previously^24^, we did not find strong evidence of polyclonality underlying tumor heterogeneity in the current study. Instead, the MDS formed the transcriptomic basis for spatial intratumoral heterogeneity of merlin immunoreactivity.^21^

In this study, we harnessed a multi-omic analysis to study whether VS^nf2^ and VS^spo^ have differences in molecular pathogenesis.^58,59^ We did not find transcriptomic, or protein expression level differences in the Schwann cell populations of VS^nf2^ and VS^spo^. Instead, both VS^nf2^ and VS^spo^ demonstrated plastic Schwann cell states along the MDS spectrum. Similar to prior studies^31,38,39^, Schwann cells could be sub-classified based on their transcriptomic phenotypes; sub-classes we found to be strongly related to their MDS status. Furthermore, a robust lineage inference strategy revealed the relatedness, and potentially plastic Schwann cell states that were driven by TEAD1 transcription factor activity.

We validated the centrality of TEAD1 to MDS, and ultimately to the mechanisms driving VS tumor progression with *in-silico*, *in vivo*, and *ex vivo* analyses. These results allow us to infer the potential utility of TEAD1 inhibition^7,50,51,60^ as a targeted therapeutic strategy in VS^nf2^ and VS^spo^, as well as potentially other *NF2*/merlin driven schwannomas. We believe that targeting TEAD1 offers tissue-specificity for Schwann cell-derived tumors since TEAD1 is a lineage-specific transcription factor for Schwann cells. ^9,10^ Our results also offer insight into the available therapeutic window with pharmaceutical TEAD1 inhibition. Recent Phase1/2 clinical data from the use of VT3989 attests to the safety in human populations.^8^ We found that TEAD auto-palmitoylation inhibition with TEADi was selective against human-derived schwannoma cells, while sparing normal Schwann cells (**Fig. 5G**). We found that despite a strong predilection for TEAD1 expression in normal Schwann cells, TEAD1 did not interact with YAP/TAZ complex, a phenomenon readily observed in the schwannoma cells (**Fig. 5E**). Exposure to TEADi prevented TEAD1-YAP/TAZ interaction and inhibited the tumor promoting signals along at least three axes (Schwann proliferation, macrophage chemotaxis/activation, and angiogenesis). TEAD1 inhibition potentially reverses Schwann cell phenotype from a high to a low MDS state and may prevent VS tumor escape seen when targeting the angiogenic pathways^61–63^ or MEK inhibition alone.^64^

## Limitations

We did not perform a detailed survey of the clonal genetic evolution of Schwann cells in VS tumors with single-cell resolution DNA sequencing. However, given the low mutational burden of VS tumors^16^, we suspect that the Schwann cell variation may be driven by epigenetic factors, partly by changes in chromatin accessibility. Currently, the safety, and long-term tolerability of TEAD1 inhibition is lacking in human patients with benign tumors such as VS. We are now creating a clinical trial to test the efficacy of TEAD1 inhibition in human patients with both VS^nf2^ and VS^spo^ tumors.

## Conclusions

We found that in VS^nf2^ and VS^spo^, tumor-driving merlin-depleted Schwann (Schwann^mer-^) cells exhibited elevated hippo activity that was predominantly driven by TEAD1. *In-silico* TEAD1 perturbation led to a reversal to merlin-intact Schwann phenotype. These findings, and tumor cell growth suppression were confirmed in NF2^fl/fl;Peri-Cre^ mouse model^7^, and in human-derived schwannoma cells with a TEAD auto-palmitoylation inhibitors.^8^ Our computational, and experimental results confirm that TEAD1 inhibition could be a potent, targeted strategy for schwannomas, and other NF2-related tumors.

## Supporting information

Supplemental Tables and Key Resources Table

## Significance

Vestibular schwannomas can lead to hearing loss and brainstem compression for which effective, targeted non-interventional strategies are lacking. The authors found sub-populations of Schwann cells have variable merlin activity depletion, and this drove tumor progression. Pharmacological TEAD1 inhibition resulted in alleviation of merlin depletion signaling, and in anti-tumor activity in NF2/merlin depleted tumors. This study forms the basis for attempting TEAD1 inhibition in patients with NF2 syndrome.

## TOC blurb

Variable merlin depletion in Schwann cells drives tumor progression in vestibular schwannomas that is targetable with TEAD1 auto palmitoylation inhibition.

## Author contributions

Authors MTL, DM, NRama, DA, SS, IR, LL, and SV generated primary data. Authors HNK and PC performed surgical approaches, and human tissue procurement. Authors LL and DP performed animal studies. Authors MTL, NRama, DB, IR, XZ, and DM generated imaging data. Authors NRama, HO, RH generated high-resolution spatial transcriptomic datasets. Author NRavi performed whole genome sequencing for NF2 patients. Authors ZA and KA generated bulk DNA methylation datasets. Author TT provided key pharmaceutical agents, and insights for study design. Authors AE, BS, and KJ generated bioinformatics datasets. Authors MTL, DB, NR, SS, IR, DM, XZ, JFS, and PC performed data analysis. Authors JS, RPH, DP, and PC provided study oversight. Authors MTL, DB, and PC generated manuscript drafts.

## Acknowledgements

The authors are clinicians dedicated to patient care. The authors thank the patients and their families for participating in the clinical trials that made this work possible.

## Methods

### Experimental model and subject details

#### Human subjects

Annotated surgical samples were obtained from non-consecutive patients (n = 15) that underwent surgery by authors HNM and PC between 2014 – 2022 (https://clinicaltrials.gov identifier NCT00060541) (**Sup. Table S1**). The study was conducted at the National Institutes of Health (NIH) Clinical Center in Bethesda, MD and approved by the Combined Neuroscience Institutional Review Board (IRB) of the National Institute of Neurological Disorders and Stroke, Bethesda, MD. Written informed consent was obtained from each patient for research study participation, and the study was conducted according to the standards set by the IRB. All patients underwent a suboccipital retro-sigmoid approach for microscope-assisted tumor resection. Vestibular schwannomas were removed in piecemeal fashion and sent for histopathological examination following formalin fixation and paraffin embedding (FFPE). Research specimens were flash frozen for *in-vitro* assays.

#### Animal subjects

Periostin-CRE mice were provided by S. Conway (Indiana University) and as previously described ^7^, mice were crossed with NF2^fl/fl^ mice (RIKEN Bioresource Research Centre) to generate Periostin-CRE;NF2^fl/fl^ mice. Age-matched CRE-littermates were used as experimental controls. All experiments included male and female animals in approximately equal numbers. Mice were housed in specific pathogen-free conditions and fed *ad lib* with standard rodent diet and water. Mice were kept no longer than 9 months to avoid substantial mortality observed in NF2^fl/fl^ mice after this time point. 5-month-old mice were treated with vehicle control or 30mg/kg of TEAD auto-palmitoylation inhibitor (VT3989, Vivace Therapeutics, San Mateo, CA) for 21 days by oral gavage. Mice were killed using carbon dioxide and cervical dislocation. Tissues were fixed in 4% paraformaldehyde, and dorsal root ganglia were dissected for further analysis. Animal experiments were approved by the Plymouth University Animal Welfare and Ethical Review Board, and conformed to UK Home Office regulations under the Animals (Scientific Procedures) Act 1986 following ARRIVE guidelines.

### Method details

*Key resources table:* Sup. Table S10.

#### Single cell RNA sequencing

Tumor samples were stored and transported in 0.9% NaCl or RPMI 1640 on ice. Samples were chopped into ∼2-4mm^3^ sized pieces and dissociated using the Human Tumor Dissociation Kit in gentleMACS C tubes using the gentleMACS dissociator (Bergisch Gladbach, North Rhine-Westphalia, Germany). Samples were briefly centrifuged at 300 rpm for 30 seconds and filtered through a 70-nm strainer. The single cell suspensions were washed with 10 mL RPMI 1640 and re-centrifuged at 2100 rpm for 5 minutes at 4°C. Single-cell suspensions then underwent red blood cell (RBC) lysis using the RBC Lysis Solution 10X (Bergisch Gladbach, North Rhine-Westphalia, Germany). Non-viable cells were removed using a Dead Cell Removal Kit (Bergisch Gladbach, North Rhine-Westphalia, Germany). Single cell suspension quality, number and viability were assessed with a dual fluorescence cell counter Luna-FL (Logos Biosystems, Gyeonggi-do, South Korea).

A total of 8,000-10,000 cells from each tumor cell suspension were washed twice with PBS+0.04% BSA and resuspended to a final concentration of 1000 cells per mL. 10X Genomics’ Chromium instrument and Single Cell 3′ Reagent kit (V3 and 3.1) were used to prepare individually barcoded scRNAseq libraries following the manufacturer’s protocol. Library quality was assessed by BioAnalyzer traces (Agilent BioAnalyzer High Sensitivity Kit) and quantified using the Qubit system. Sequencing was performed on the Illumina NextSeq machine, using the 150-cycle High Output kit with 28 bp read 1, 8 bp sample index, and 91 bp read 2. Following sequencing, bcl files were demultiplexed into FASTQ, aligned to human transcriptome GRCh38, and single-cell 3′ gene counting was performed using the standard 10X Genomics’s CellRanger mkfastq software (V3.0.2).

#### Single nuclei multiome sequencing

Nuclei were isolated from frozen tissues using an adaptation of a previously established protocol^65^. Samples were washed with 800uL of Detergent-Lysis Buffer (0.1% Triton-X). Tissues were chopped into small pieces, homogenized in a 7mL Dounce Homogenizer (Fisher Scientific # 501945204) on ice, and passed through a 70 mm MACS strainer (Miltenyi Biotech #130-098-462). Nuclei were spun down at 3200 xg for 5 minutes at 4°C and processed using the Chromium nuclei isolation kit (10x Genomics #1000494). Briefly, nuclei were resuspended in 1mL of low sucrose buffer, and gently layered above 4 mL of high sucrose buffer taking care not to disrupt the density gradient. The gradient was spun down at 3200 xg for 20 minutes at 4°C, then resuspended in Nuclei Resuspension Buffer. Aliquots of resuspended nuclei were stained with Acridine Orange dye (Logos Biosystems # F23001) and counted using an automated fluorescence cell counter (Logos Biosystems # L20001).

Nuclei were immediately processed using a Single Cell Multiome ATAC + Gene Expression Assay (10X Genomics, # 1000285) following manufacturer recommendations. Nuclei were loaded onto a Next GEM Chip J (10X Genomics # 1000230), targeting a yield of 3,000 nuclei. Library preparation was performed according to the manufacturer’s recommendations. Libraries were pooled and sequenced on an Illumina NextSeq 500 machine using a pair of High-Output Reagent Kits v2.5 with 150 cycles (Illumina # 20024907). Sequencing reads were demultiplexed and aligned to the hg38 reference genome using the 10X Genomics Cell Ranger software (arc v2.0.1) mkfastq function with default settings, and counts were generated using the Cell Ranger-arc count function.

#### Single cell/nuclei RNA and multiome analysis

Filtered and raw .h5 files from Cell Ranger were processed in RStudio using Seurat v5.0.1.^66^ Cell Free RNA contamination and doublets was estimated and removed using the SoupX^67^ and DoubletFinder^68^ R packages. For quality control, nuclei and cells exhibiting high mitochondrial RNA content (>25%), low transcript complexity (fewer than 300 detected genes; nFeature_RNA < 300), or low total read counts (nCount_RNA < 500) were excluded, yielding 77,930 nuclei and 15,500 cells after filtering. The data was log normalized using NormalizeData(), and variable features were identified using FindVariableFeatures(). The data was scaled using ScaleData() before principal component analysis using RunPCA(). Single-nuclei samples underwent marker-based classification using scSorter^69^, referencing Schwann cells and associated schwannoma microenvironment canonical cell markers. Cell-based classification was validated by manually comparing predictions from scSorter with clusters identified by uniform manifold approximation and projection (UMAP) and unsupervised clustering functions: RunUMAP(dims=1:30), FindNeighbors(dims = 1:30), FindClusters(resolution = 0.5). Differentially expressed genes (DEGs) were identified for each cell class using the Seurat FindAllMarkers() function with the default settings (Log2-fold change >0.25, and gene must be expressed in >25% of cells in that cluster). S and G2M cell cycle phase scores were determined using the CellCycleScoring() function in Seurat.

Copy number variants (CNVs) were first identified using the R-packages InferCNV^70^ with endothelial cells, macrophages, neutrophils, and pericytes marked as reference cells. In parallel, CONICSmat^71^ was used to produce UMAP plots that describe inferred chromosomal expression. Chromosome 22q loss was determined using the inflection point of the chromosome 22q expression density plot in samples that had adequate loss in Schwann cells as visualized using the FeaturePlot() function (n= 7). ‘The common upregulated genes among the single-nucleus samples (n=6) of the cells denoted as having a loss in the 22q chromosome with an average fold change of greater than 0 and an adjusted p-value less than 0.05 were used to create a merlin depletion score (MDS) using the tl.rank_genes_groups() function in Scanpy.^72^ MDS class was determined for Schwann cells based merlin depletion score quartiles: +σ > 2σ > -σ as merlin-depleted (Schwann^mer-^), merlin-intermediate (Schwann^mer+/-^), and merlin-intact (Schwann^mer+^). Similar module scoring was done using DEGs from NF2-depleted cell-lines for validation and for genes involved in the hippo pathway. For additional validation, normal Schwann cells were sourced from 2 Normal spinal cord public datasets (GSE190444, n = 8 from 7 donors; GSE243077, n = 9)^32,33^ and one normal auricular nerve dataset (GSE230375, n = 2)^18^ on NCBI-GEO. All samples were preprocessed using the same pipeline described above. Normal Schwann cells (n = 2,395) were then combined with VS tumors’ Schwann cells, and the merlin depletion score was recalculated.

Individual datasets were integrated using the CCA integration method from the Seurat v4^73^ R-package integration workflow to remove batch effects while preserving biological variation. Integration anchors were found using FindIntegationAnchors() with the number of dimensions set to 30, and each sample’s gene expression matrix was integrated into one object via IntegrateData(). Integrated objects were scaled, run through a PCA analysis, and clustered through Uniform Manifold Approximation and Projection (UMAP). The Schwann cells and macrophages were isolated from the integrated object and were processed through the standard Seurat pipeline. Unsupervised clustering of each subset of the integrated object were manually labelled after cross-referencing with existing literature using the FindAllMarkers() function and based on gene-set enrichment analysis (GSEA) using the clusterProfiler^74^ R-package and GSEApy^75^ python package. Cell-cell communication networks were identified using the CellChat^76^ R package.

snRNAseq data was visualized using the Scanpy python package. To assess sample-level transcriptomic similarity of cell types between VS^nf2^ and VS^spo^, cosine similarity distances between all sample–cell type pairs were computed using the tools.Distance() function in Pertpy.^77^ A Mann–Whitney U test was then applied to evaluate differences in transcriptomic similarity within versus between VS^nf2^ and VS^spo^ samples across cell types. For RNA velocity and lineage tracing analysis, the python package scvelo^78^ was used to calculate RNA velocities using the unfiltered expression matrix from 10x Cell Ranger software, and ratios of spliced to unspliced mRNA calculated from velocyto.^44^ The CellRank^45^ python package was used to compute a transition matrix, initial and terminal states, cell lineage fate probabilities, and lineage specific gene drivers. Partition-based graph abstraction (PAGA)^79^ was used to infer directional graph based on RNA velocity.

To assess global chromatin accessibility, the ATACseq reads and fragments for each sample were integrated together and preprocessed for quality control using ArchR^80^ with peaks identified using MAC2^81^ for 36,129 cells. The ATAC ArchR project was subset for Schwann cells and BigWig files were generated for each MDS class. Using deepTools^82^, the count matrices of peaks identified by MACS2 and gene regions (TSS-1000bp to TES+1000bp) were computed for each BigWig files and were visualized using plotHeatmap(). pyCirclize^83^ was used to plot the relative fragment coverage for Schwann^mer-^, and Schwann^mer+^ across the entire genome (bin size = 2.5 million base pairs) and Wilcoxon test was used to determine statistical significance. Individual gene coverage plot was generated using a custom python function.

For transcription factor activity, preprocessed chromatin accessibility profiles from ArchR were analyzed using pycisTopic to perform topic modeling, grouping peaks into co-accessible topics. Peaks were annotated with transcription factor motifs, and per-cell topic distributions were obtained. For 17,154 cells with matched snATAC-seq and snRNA-seq data, SCENIC+^48^ was used to integrate chromatin accessibility with gene expression, linking accessible motif-enriched peaks to nearby genes. Gene regulatory networks were constructed and refined by motif enrichment, peak accessibility, and correlation with target gene expression. Motif enrichment was performed using the hg38 cisTarget databases (hg38_screen_v10_clust) together with motif-to-gene annotations (motifs-v10nr_clust-nr.hgnc). Finally, gene regulon activity was quantified in each cell using enrichment-based scoring, yielding cell-level transcription factor activity profiles. SCENIC+ was also used to visualize the top five transcription factors in each stromal cell type and Schwann MDS class. Selected transcription factor activities in Schwann cells were further scaled and smoothed across the merlin depletion score using pyGAM^84^.

Perturbation analysis was performed using SCENIC+ to evaluate the regulatory influence of TEAD1. The perturbation was simulated by setting TEAD1 activity to 0, and propagating the effect through the inferred regulatory network over five iterations. Significant changes in gene expression and MDS before and after perturbation were assessed using a Wilcoxon rank-sum test.

#### Chromatin immunoprecipitation-sequencing (ChIPseq)

Surgical tissues were fragmented into 2-4 mm^3^ pieces and cross linked using 1% paraformaldehyde (Sigma #F8775) for 10 minutes, followed by glycine quench for 5 minutes at room temperature. ChIP was performed using an EZ-Magna ChIP kit (Millipore Sigma #17-10086) following manufacturer’s instructions. Sonication was optimized for the tissue type, and fragment sizes ∼ 200kb were verified using gel electrophoresis. Sonication was conducted on wet ice using 10 cycles of 10 seconds on, 50 seconds off at 75% power (Qsonica #Q125-110). Immunoprecipitation was performed overnight at 4 °C on a rotating rocker using 10 mL of specific antibodies. After elution and reversal of cross-linking, DNA fragments were purified, and libraries were prepared using the Illumina TruSeq ChIP Library Preparation Kit following manufacturer’s instructions (Illumina # IP-202-1012). Sequencing was performed on an Illumina NextSeq machine using the NextSeq 500/550 Mid Output Kit v2.5 with 150 Cycles (Illumina # 20024904).

Data was analyzed using the Galaxy ChIP-seq pipeline. Briefly, raw QC reads were filtered using adapter trimming program Trimmomatic, followed by alignment using BWA. Next, normalized bigwig files were created and scaled using BEDTools.^85^ DeepTools was used to generate gene-body meta-profiles from bigWig files and to calculate genome-wide IP enrichment. Peaks were annotated relative to gene features using MACS2. MACS2 peaks were further analyzed in R using ChIPseeker^86^ and GenomicRanges.^87^

#### DNA methylation

Tissue sections (5 μm) were cut from formalin-fixed paraffin-embedded surgically derived vestibular schwannoma specimens. DNA was isolated and purified (Qiagen DNeasy #69556). After sodium bisulfite conversion using the Zymogen EZ DNA Methylation kit (#D5001), 1ug each of methylated and unmethylated DNA was analyzed using the Illumina Infinium MethylationEPIC chip (#20087706). Samples were randomly assigned to plates. Signal intensities were retrieved and poorly performing probes were removed (P value > 0.01 for > 1% CpGs, ch probes, normal rs probes, CpGs on sex chromosomes, SNP probes, cross-reactive probes, invariable probes). Data was processed and analyzed using the MissMethyl R package.^88^ Beta values were calculated for each probe and adjusted for cell type composition. Probes associated with genes of interest were identified and compared across samples. P values were adjusted for multiple comparisons using false discovery rate analysis with Benjamini-Hockberg statistics. Cluster Heatmap and UMAP visualizing beta values for bulk DNA methylation of CNS tumors were created using standard Python packages.

#### Germline genotyping

Germline genetic testing was performed by whole genome sequencing (WGS) of peripheral blood mononuclear cell pellets in NF2 patients. Briefly, DNA was extracted using the Qiagen AllPrep DNA/RNA extraction kit (Catalog no: 80204, Qiagen, USA) and quantified on the Qubit 4 fluorometer (ThermoFisher Scientific, USA). Genomic DNA (500 ng) was used to construct sequencing-ready libraries with the NEBNext Ultra II DNA Library Prep Kit for Illumina (E7645) and NEBNext Multiplex Oligos for Illumina (E7395) (New England Biolabs, USA). Libraries were sequenced on the Illumina NovaSeq X instrument (Illumina, USA) and resulting FASTQ files were processed using the NF-core Sarek pipeline to call single nucleotide and copy number variants.^89^ Variant annotation was performed using ANNOVAR (version 2020-06-08) and snpEff (version 5.2).^90,91^ High impact loss-of-function variants affecting coding and splice site regions of the canonical NF2 RefSeq transcript (NM_000268.4) were retained for further analysis and classified according to the American College of Medical Genetics and Genomics and Association for Molecular Pathology guidelines.^92^

#### Multiplex immunohistochemistry

Multiplex immunohistochemistry (mIHC) was performed on 5 µm-thick paraffin-embedded sections. Sections were deparaffinized and treated using a standard antigen unmasking step in 10 mM Tris/EDTA buffer pH 9.0. Sections were then blocked with Human BD Fc Blocking solution (BD Biosciences #564219) and treated with TrueBlack Reagent (Biotium #23014-T) to quench intrinsic tissue autofluorescence. The sections were then immunoreacted for 1 hour at RT using 1 µg/ml cocktail mixture of immunocompatible antibodies. Primary antibodies were either directly conjugated or indirectly labelled with secondary antibodies using the following spectrally compatible fluorophores: Alexa Fluor 430, Alexa Fluor 488, Alexa Fluor 546, Alexa Fluor 594, Alexa Fluor 647, IRDye 800CW. After washing off excess antibodies, sections were counterstained using 1 µg/ml DAPI (Thermo Fisher Scientific #D1306) for visualization of cell nuclei. Slides were then mounted using Immu-Mount medium (Thermo Fisher Scientific #FIS9990402) and imaged using an Axio Imager.Z2 slide scanning epifluorescence microscope (Zeiss) equipped with a 20X/0.8 Plan-Apochromat (Phase-2) non-immersion objective (Zeiss), a high-resolution ORCA-Flash4.0 sCMOS digital camera (Hamamatsu), a 200W X-Cite 200DC broad band lamp source (Excelitas Technologies), and 7 customized filter sets (Semrock). Image tiles (600 x 600 µm viewing area) were individually captured at 0.325 micron/pixel spatial resolution, and tiles were seamlessly stitched into whole specimen images using the ZEN 2 image acquisition and analysis software program (Zeiss). Pseudocolored stitched images were exported to Adobe Photoshop and overlaid as individual layers to create multicolored merged composites.

For quantitative analysis, raw TIFF images from mIHC were downsampled using the Lempel–Ziv–Welch lossless compression algorithm. Nuclear segmentation was performed in QuPath^93^ using the Cell Detection module on the DAPI channel (Background radius = 0; Median filter radius = 0; Sigma = 3; Minimum area = 10 px; Maximum area = 1000 px; Threshold = 5; Cell expansion = 20). A total of 2,721,272 cells were identified across all tumor samples. Random Forest classifiers were trained to determine marker positivity and annotate cells. Anndata objects were constructed from exported measurements, and the Leiden algorithm was applied to cluster marker-positive cells into cell types. VEGFA⁺ cells were defined by thresholding at the mean expression value.

To investigate how hypoxic stress may lead to angiogenesis, vessels were annotated by creating spatial clusters of endothelial cells using DBSCAN^94^, computing convex hull of cluster, and then thresholding the circularity of this structure to ensure all vessels used for analysis were of regular and comparable shapes (circularity >= 0.7). Normalized distances between Schwann VEGFA + cells and Vessels were then calculated relative to the mean intercellular distance within each sample.

#### RNAscope in situ hybridization

mRNA was visualized using RNAscope *in situ* hybridization (ACDBio, USA) following the manufacturer’s instructions^95^. Briefly, probes were warmed for 10 minutes at 40°C in a water bath, then cooled to room temperature. Probes C2, C3 and C1 were mixed in a 1:1:50 ratio. 5 µm-thick paraffin-embedded tumor sections were deparaffinized and placed in HybEZ Slide Racks. Probes were hybridized to target mRNAs by incubating probe mixtures in a HybEZ oven using a HybEZ Humidity Control Tray for 2 hours at 40 °C. Slides were washed twice for 2 minutes at room temperature using 1X Wash Buffer. Slides were sequentially incubated in AMP 1-FL for 30 minutes at 40 °C using the HybEZ oven, AMP 2-FL for 15 minutes at 40 °C, AMP 3-FL for 30 minutes at 40 °C, and AMP 4-FL for 15 minutes at 40 °C, with two 1X Wash Buffer washes between incubation steps. Slides were counterstained with DAPI for 30 seconds at room temperature, then mounted using fluorescent mounting medium.

Slides were imaged using an Axio Imager.Z2 slide scanning epifluorescence microscope (Zeiss) equipped with a 20X/0.8 Plan-Apochromat (Phase-2) non-immersion objective (Zeiss), a high-resolution ORCA-Flash4.0 sCMOS digital camera (Hamamatsu), a 200W X-Cite 200DC broad band lamp source (Excelitas Technologies), and customized filter sets (Semrock). Image tiles (600 x 600 µm viewing area) were individually captured at 0.325 micron/pixel spatial resolution, and tiles seamlessly stitched into whole specimen images using the ZEN 2 image acquisition and analysis software program (Zeiss). Pseudocolored stitched images were then exported to Adobe Photoshop and overlaid as individual layers to create multicolored merged composites.

For quantification, image analysis was performed as described for mIHC. Raw TIFF files were downsampled using the Lempel–Ziv–Welch compression algorithm, and nuclear segmentation was performed in QuPath (Cell Detection parameters as above). A total of 39,699 cells were identified across all tumor samples. Random Forest classifiers were trained to determine marker positivity and annotate cells. Anndata objects were generated from exported data, and the Leiden algorithm was used to cluster marker-positive cells into cell types. VEGFA⁺ cells were defined by thresholding at the mean expression value.

#### Western immunoblotting

Tissue samples were washed with ice-cold PBS and whole-cell lysates were collected on ice using a radioimmunoprecipitation assay (RIPA) lysis buffer (Thermo Scientific #89900) containing a protease inhibitor cocktail (Halt protease inhibitor, Sigma-Aldrich #539137). Whole-cell lysates were quantified using the bicinchoninic acid (BCA) protein assay (Thermo Scientific #23227). Proteins were electrophoretically separated on 4%–12% NuPAGE Bis-Tris gels (Invitrogen #NP0321BOX) and electroblotted onto polyvinylidene difluoride (PVDF) membranes with the Trans-Blot Transfer Turbo System (Bio-Rad). After blocking membranes for 1 hour in 5% bovine serum albumin (BSA) in 0.05% TBS-Tween at room temperature, they were incubated overnight at 4° C with primary antibodies. Membranes were washed with TBS-Tween and incubated with peroxidase-conjugated secondary antibody for 1 hour at room temperature while rocking. Membranes were developed with ECL (Super Signal West Pico PLUS, Thermo Fisher #34579). Immunoreactive signal was detected and imaged using the ChemiDoc MP Imaging System (Bio-Rad).

#### Xenium Spatial Transcriptomic Analysis

Formalin-fixed paraffin-embedded (FFPE) tissue sections were processed for spatial transcriptomic profiling using the Xenium In Situ Analyzer (10x Genomics) following the manufacturer’s FFPE Cell Segmentation workflow with minor optimizations to incubation times and reagent preparation. FFPE sections (10 µm) were mounted onto Xenium slides and baked at 60 °C for 2 hours to ensure adhesion and complete paraffin softening. Sections were deparaffinized through serial xylene and graded ethanol washes, followed by rehydration in 70% ethanol and water. After equilibration at room temperature, slides were assembled into Xenium cassettes. Tissue decrosslinking was performed in Xenium FFPE Tissue Enhancer containing 8 M urea and diluted Perm Enzyme B at 37 °C for 40 min on a Bio-Rad C1000 Touch Thermocycler. Following three 1-min washes in PBS + 0.05% Tween-20 (PBST), custom probes corresponding to a 300-gene panel were resuspended in TE buffer and combined with Xenium Probe Hybridization Buffer and additional TE to prepare the hybridization mix. Probe mix was briefly heated to 95 °C for 2 min to equilibrate, cooled on ice, and then added to each cassette for hybridization at room temperature for 16–18 hours on a thermomixer.

Post-hybridization washes, ligation, and rolling-circle amplification were performed sequentially using the manufacturer’s reagents (Modules A and B) and thermocycler programs, with intermediate PBST washes between each step. Amplified products were fixed and blocked in Xenium Block & Stain Buffer before incubation with the Multi-Tissue Stain mixture for nuclear segmentation, which proceeded overnight at 4 °C. Slides were treated with Xenium Stain Enhancer and Autofluorescence Reduction reagents in subdued light, followed by a graded ethanol dehydration and rehydration series. Tissues were equilibrated in PBST, stained with Xenium Nuclei Stain Buffer, and prepared for instrument loading. Two slides were run simultaneously on the Xenium Analyzer following the manufacturer’s automated imaging and chemistry protocol. Four reagent bottles (nuclease-free water, Buffers A and B, and Probe Removal Buffer containing DMSO, KCl, and Tween-20) were loaded along with fresh Modules A and B and standard Xenium consumables. Image acquisition and transcript decoding were performed automatically by the Xenium Analyzer software. After completion of the run, slides were removed, rinsed in PBST, and stored at 4 °C in the dark for subsequent H&E staining. Raw Xenium output files were exported for downstream analysis.

The raw Xenium output files for eight vestibular schwannoma (VS) samples were preprocessed, filtered for quality control (minimum of 10 counts per cell, retaining 1,219,325 cells after filtering), and integrated using Scanpy and the SpatialData^96^ packages in Python. Cell type, sub-classes, and MDS class were classified through annotation transfer from the integrated single-cell and single-nucleus RNA sequencing datasets using TACCO.^40^

To explore NF2 transcript sub-cellular localization, cellular and nuclear boundaries were delineated with the Xenium Multi-Tissue Stain Mix. Spatial transcript localization and quantification were performed in Python using GeoPandas and Shapely to assign transcript coordinates to segmented cells based on sub-cellular spatial localization. Fisher’s exact test was used to determine nuclear or cytosolic enrichment of NF2 transcripts in Schwann MDS class and stromal cells.

Neighborhood enrichment scores were computed across the integrated object using Squidpy^30^ to examine inter-cellular interaction in VS. These interactions were further explored by identifying spatial niches with CellCharter^43^ based on the scVI^97^ latent space and Squidpy-derived spatial neighborhood graph across the integrated object. The optimal number of niches was determined by cluster stability analysis, and CellCharter was further applied to quantify cell type and subclass enrichment within each niche.

To continue to investigate how hypoxic stress may lead to angiogenesis, DBSCAN clustering was used to define Schwann Hypoxic Regions (spatial clusters of hypoxic Schwann cells) and Vessel (spatial clusters of endothelial cells). The centroids of Schwann Hypoxic Regions and Vessel were correlated (Pearson’s r) and normalized distances between Schwann Hypoxic cells and Vessels were calculated relative to the mean intercellular distance within each sample. All visualizations were generated using Scanpy, Squidpy, Seaborn^98^ and Matplotlib.

#### Quantification and statistical analysis

Statistical details of experiments have been included in the figure legends. Comparative analysis was performed using paired or unpaired T tests after testing for variance equality, and using Welch’s approximation for degrees of freedom, or using one/two-way ANOVAs where appropriate. P values < 0.05 two-tailed were considered statistically significant. Statistical analyses were performed using GraphPad Prism 9.0 (GraphPad Software, La Jolla, California), or appropriate python packages.

#### Primary Cell Culture

Clinical tissue samples were collected aseptically after surgery in cold DPBS. The tissue was minced into small pieces using a sterile scalpel and suspended in cold DPBS. Contaminated blood was removed by centrifugation at 300 × g for 5 minutes under cold conditions. The tissue samples were then resuspended in 1 ml of a Trypsin-EDTA and Collagenase solution (1:1 mixture of 0.25% Trypsin-EDTA and 0.2% Collagenase) and incubated for 30 minutes in a CO₂ incubator at 37°C. To neutralize the effect of Trypsin, 5 ml of complete medium was added, followed by centrifugation at 300 × g for 5 minutes. After carefully removing the supernatant, the digested tissues were suspended in complete medium and plated onto Petri dishes and left undisturbed for 3–5 days in a CO₂ incubator at 37°C. Cells were cultured in Schwann media in presence of 10% FBS and used for experiments after 2–3 passages. We derived primary cells from patient of NF2 schwannoma - S2290, NF2 meningioma – S2348, Sporadic vestibular NF2 schwannoma - S2470. The normal Schwann cells were bought from Neuromics Inc, and all the primary cells were cultured in Schwann growth medium (SGM001).

#### Cell Viability Assays

Cultured cells were seeded in 96-well plates at a density of 10,000 cells/well and allowed to grow for 16 hours prior to treatment. Cells were then treated with increasing concentrations (0.06 µM, 0.124 µM, 0.25µM, 0.5 µM, 1 µM, 2 µM, 4µM) of compounds #1, 2, 3, and 4 for 24, 48, and 72 hours before the viability assay. Cell viability was measured using the CellTiter-Glo kit (Promega #G7571) according to the manufacturer’s instructions. After a 30-minute incubation at room temperature, luminescence intensity was recorded using a Synergy Neo2 microplate reader (BioTek).

#### Co-immunoprecipitation (IP)

Confluent cells (treated or untreaded) were harvested using 0.25% Trypsin-EDTA (Gibco #25200-056) and rinsed with ice cold PBS. Cell pellet from each plate (10 cm) was treated with 0.5 ml ice-cold 1X cell lysis (Cell signaling #9803) and incubated on ice for 10 min with mild vertexing in between. centrifuged for 10 min at 4°C, 14,000 x g and the supernatant were collected to a new tube. The protein was estimated using standard BCA method and stored at -800°C for further use of WB or CoIP. Following the Cell signaling Immunoprecipitation protocol Pan-TEAD Primary antibody (#13295) at a dilution of 1:100 was allowed to bind overnight with rotation to form the immunocomplex. The immunocomplex was allowed to bind further with equilibrated magnetic beads (#73778) for one hr at room temperature with rotation followed by five times wash with 500 μl of 1X cell lysis buffer. Pan-TEAD protein or complex was eluted with 3X SDS sample buffer after heating the sample at 95-100°C for 5 min. Samples were analyzed by western blot

#### Western Blot Analysis

Cell lysates containing 20 µg of protein were used for sample preparation. Proteins were electrophoretically separated on 4%–12% NuPAGE Bis-Tris gels (Invitrogen #NP0321BOX) and transferred onto polyvinylidene difluoride (PVDF) membranes using the Trans-Blot Transfer Turbo System (Bio-Rad). Membranes were blocked for 1 hour at room temperature in 5% bovine serum albumin (BSA) prepared in 0.05% TBS-Tween. Following blocking, membranes were incubated overnight at 4°C with primary antibodies. After incubation, membranes were washed with TBS-Tween and then incubated with a peroxidase-conjugated secondary antibody for 1 hour at room temperature with gentle rocking. Protein bands were detected using enhanced chemiluminescence (ECL) reagent (SuperSignal West Pico PLUS, Thermo Fisher #34579). Immunoreactive signals were visualized and imaged using the ChemiDoc MP Imaging System (Bio-Rad). The antibodies used in this study were primarily from Cell Signaling Technology, including anti-pan-TEAD (S13295), anti-YAP (4912S), anti-TAZ (83669S), anti-VEGF-A (50661S), anti-rabbit (7074S), and anti-mouse (7076S). Additional antibodies were obtained from other suppliers: Thermo Fisher (anti-MCSF, 21113), Origene (anti-IL34, TA811728), and Santa Cruz Biotechnology (anti-NRG1, sc-393006; anti-Vinculin, sc-59803; anti-GAPDH, sc-47724). For immunoprecipitation (IP) samples, we used the Mouse Anti-Rabbit IgG HRP-conjugated secondary antibody (#5127) to avoid masking caused by denatured IgG heavy chains (∼50 kDa).

## Supplemental Figures

**Sup. Fig. S1:**
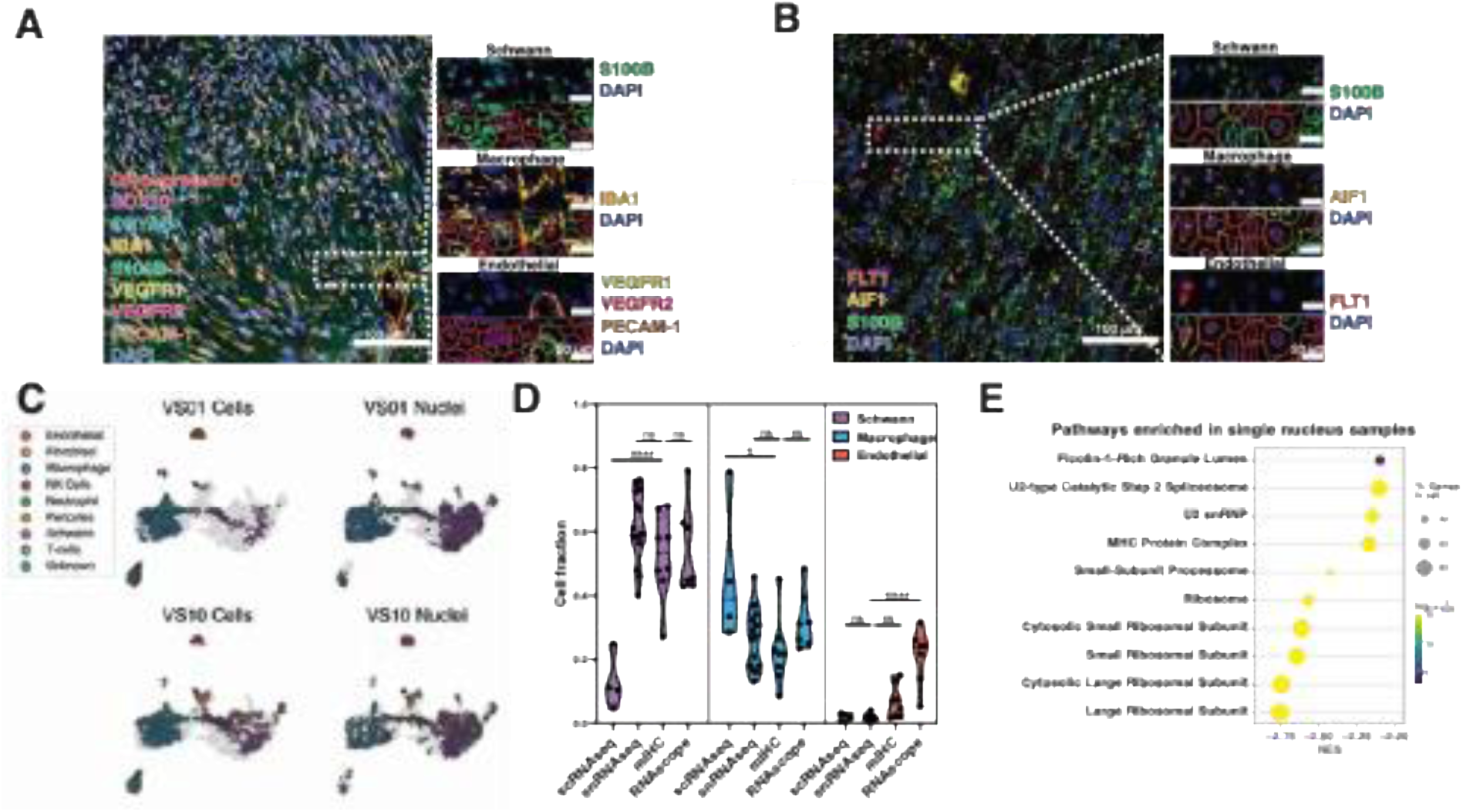
Cell classification strategy and assay comparison. **A.** Representative merged multiplexed immunohistochemistry (mIHC) image of VS01 (scale bar = 100µm). The hues of protein targets are represented by hues of the respective sub-titles. Inset is enlarged to demonstrate QuPath-based expression analysis, segmentation and cell class identification (scale bar = 20µm). **B.** Representative RNAscope in-situ hybridization, and QuPath analysis as in A. **C.** Representative tumor sample UMAP of VS01 (top) and VS10 (bottom) of canonical cell class distribution between scRNAseq (left), and snRNAseq (right). **D.** Violin plot representing the cell fractions for Schwann cells, macrophages and endothelial cells per assay method. Each point represents a single tumor tested. (ANOVA with Dunnett correction for multiple comparisons, p < 0.5 *, p < 0.0001 ****). **E.** Pairwise geneset enrichment analysis (Hallmark 2024) for pathways enriched in single nucleus RNAseq samples compared to single cell RNAseq assays.

**Sup. Fig. S2:**
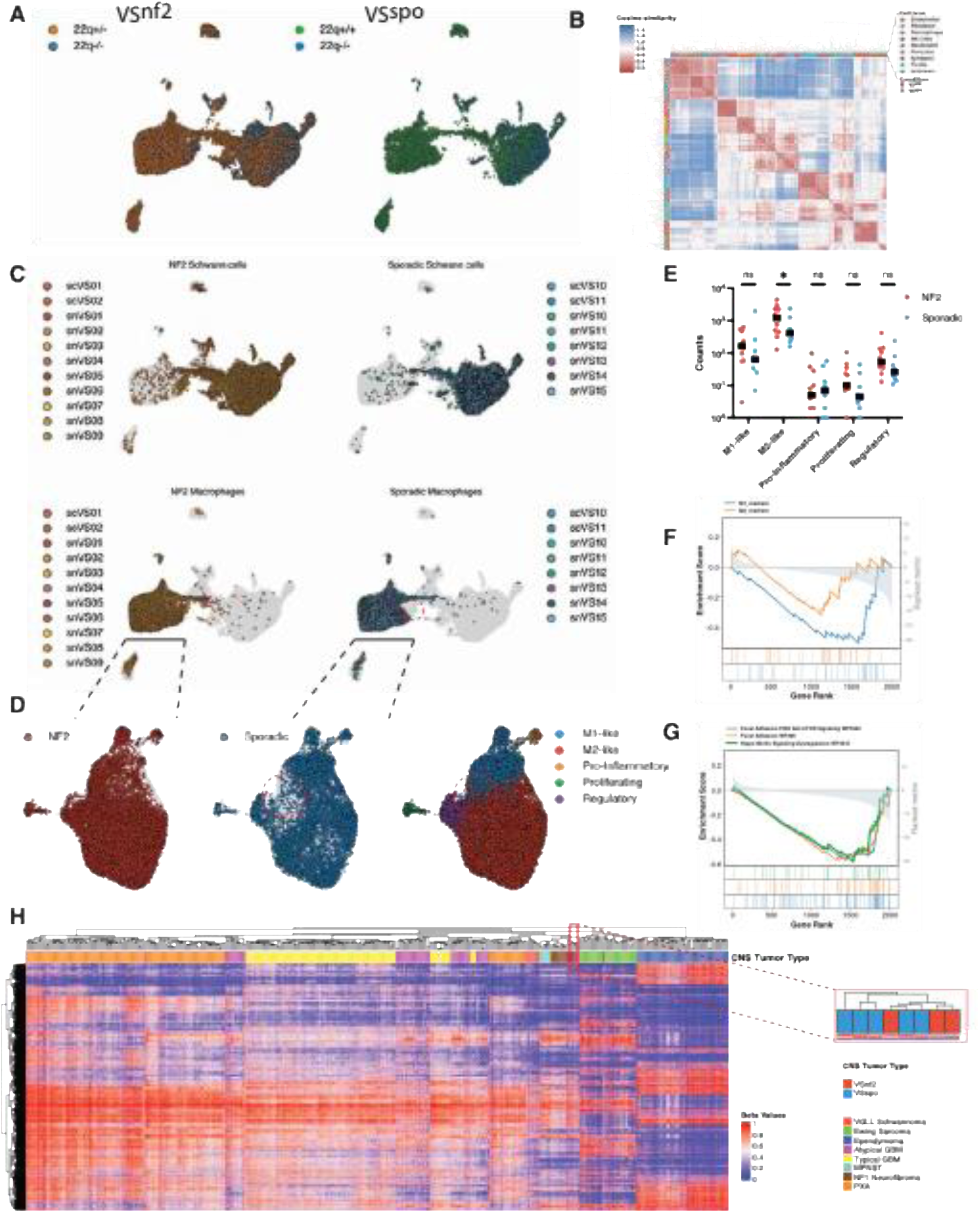
Comparison of NF2 related (VS^nf2^) and sporadic (VS^spo^) vestibular schwannoma. **A.** Uniform manifold approximation and projection (UMAP) of integrated single nucleus RNAseq (snRNAseq) and single cell (scRNAseq) datasets. Datasets were subset according to the germline condition (VSnf2 vs. VSspo) and mapped for cellular NF2 dosage following inferred copy number variant analysis for chromosome 22q. **B.** Heatmap of transcriptomic cosine similarity distance of cells with canonical cell classes, and germline condition marked. **C.** Concurrent UMAPs demonstrating the distribution of Schwann cells (top), and macrophages (bottom) from VS^nf2^ (left) and VS^spo^ (right). Hues represent the source tumors. The complete, integrated dataset is plotted as a gray background. **D.** UMAP of macrophages from VS^nf2^ (left), and VS^spo^ (middle) mapped over the total macrophage dataset as gray background. Macrophage sub-classes mapped over the total macrophage dataset with hues as represented in the legend. **E.** Dot plot demonstrating the macrophage sub-class counts across tumors. Each point represents a tumor (ANOVA with Dunnett correction for multiple comparisons, p < 0.5 *). **F.** Gene set enrichment (GSEA) plot for macrophages derived from VSnf2 vs. VSspo mapped with a macrophage state specific geneset (from Xue et al. 2014, Immunity. PMID: 24530056). **G.** GSEA plot as in F. using Wiki Pathways 2024 Human. **H.** Clustered heatmap of bulk DNA methylation beta values summarizing the differentially methylated regions for CNS tumors. Inset magnifies the clustering of VS^nf2^ and VS^spo^ in the heatmap.

**Sup. Fig. S3:**
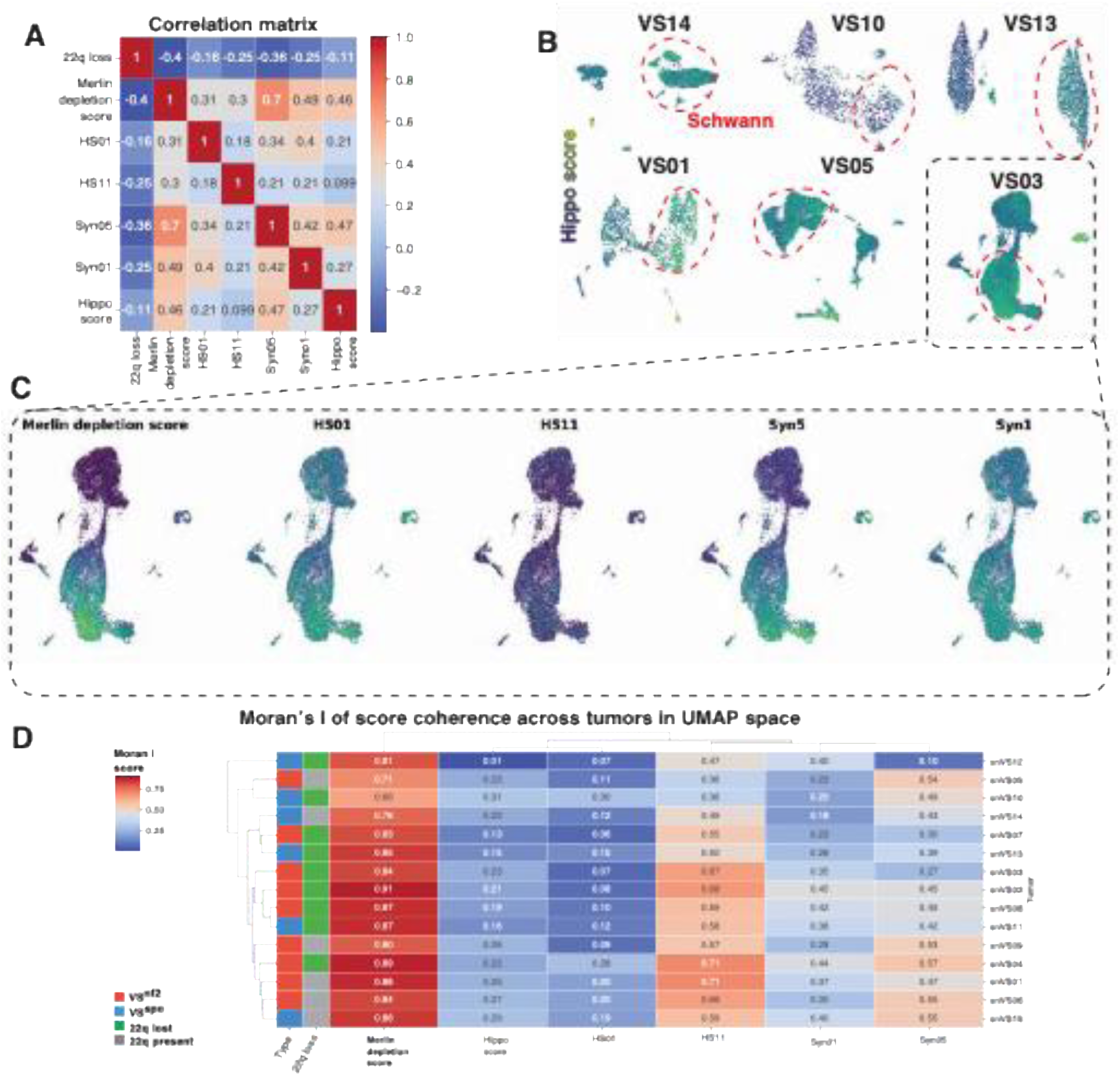
Merlin depletion score (MDS) is a discriminative and biologically coherent measure of merlin functional loss. **A.** A correlation matrix (Pearson correlation coefficient) of the merlin depletion score, chromosome 22q loss signal from copy number variant analysis, and gene expression scores derived from experimental NF2 deletion in cell lines (from Synodos for NF2 Consortium et al. 2018, PloS One; PMID: 5129897904). **B.** Representative uniform manifold approximation and projection (UMAP) of individual tumors as in Fig. 2A, with hippo gene activation score (from Wang et al. 2018, Cell Rep; PMID: 30380420) overlaid. **C.** UMAPs of VS03 tumor with scores from differentially expressed genes (DEGs) in *NF2*-depleted cell-lines, and merlin depletion score overlaid. **D.** Moran’s I score for autocorrelation of scores over the UMAP embedding. Rows represent tumors, and columns represent the scores tested for Moran’s I autocorrelation.

**Sup. Fig. S4:**
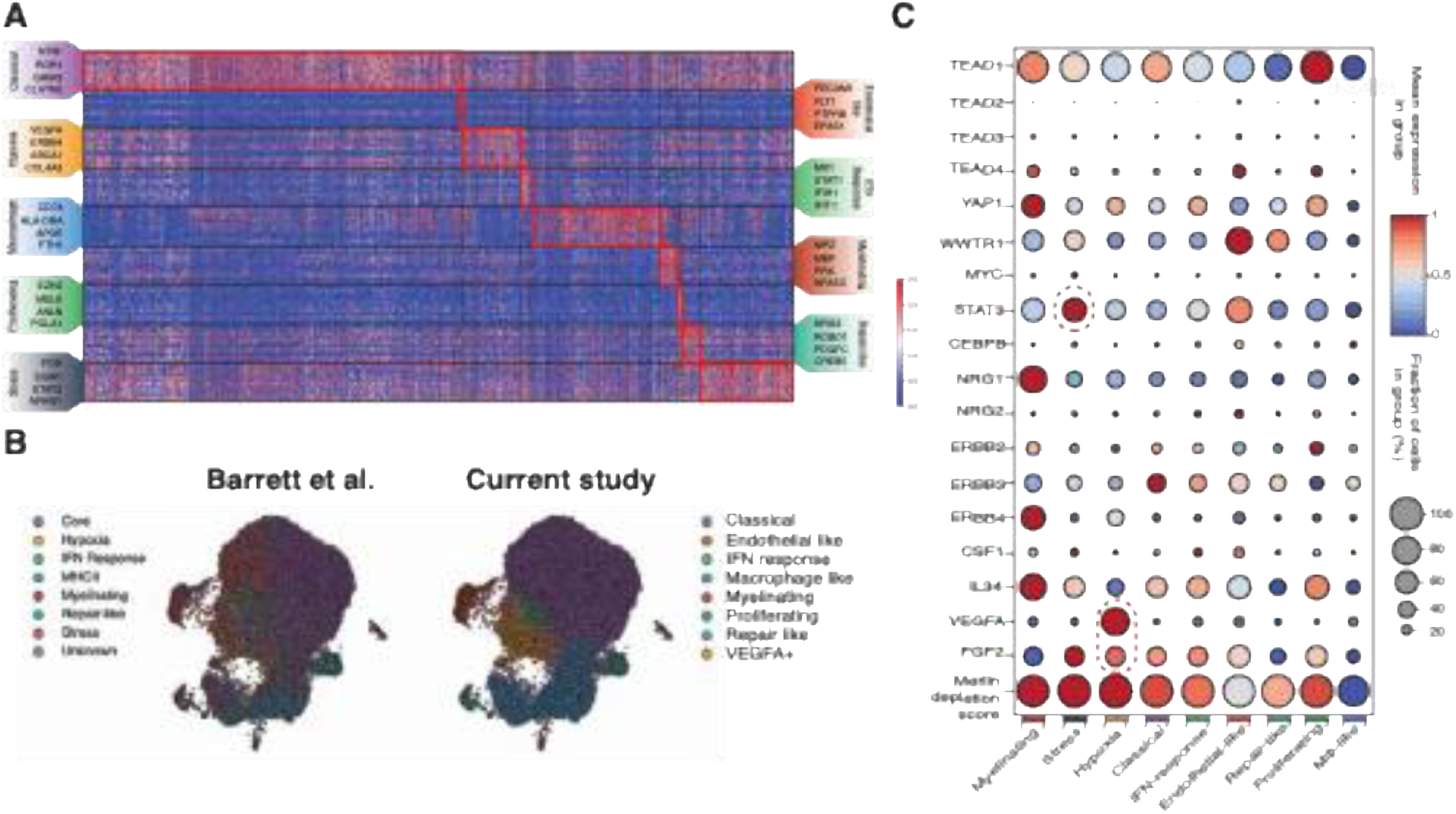
Schwann cell sub-classification schema. **A.** A clustered heatmap of top 25 differentially expressed genes among Schwann cells. Each row represents a Schwann cells, and each row cluster represents a sub-class. Over-expressed genes are outlined with red rectangles, and key genes are highlighted in boxes adjacent to each of the row cluster. **B.** A side-by-side comparison of previously published Schwann cells sub-classes and the current study, overlaid on a uniform manifold approximation and projection (UMAP) of Schwann cells from the current study. **C.** Dot plot showing the averaged gene expression profiles and merlin depletion scores of Schwann cell sub-classes. Key genes are highlighted with red ellipses.

**Sup. Fig. S5:**
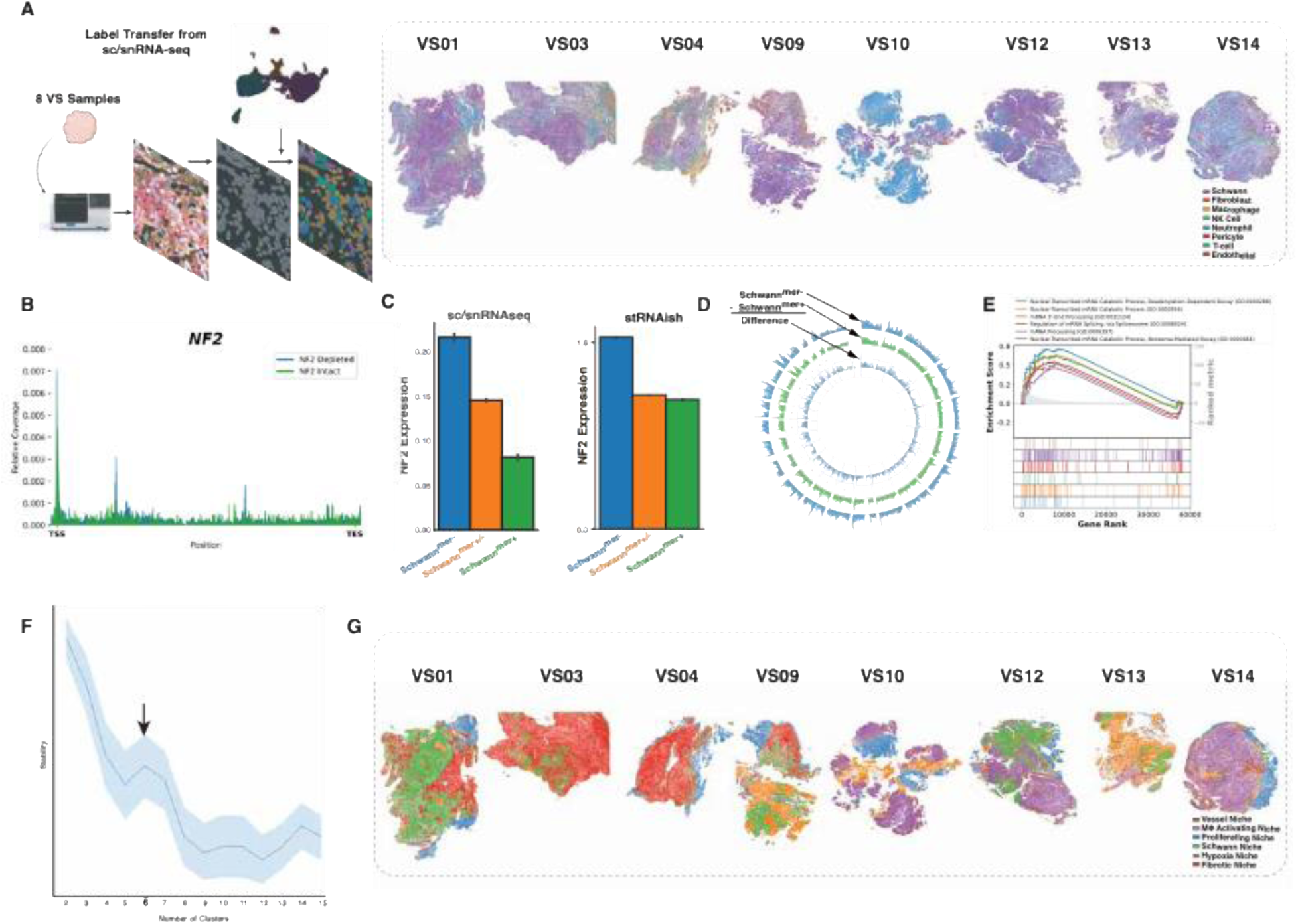
Schwann cell merlin depletion score (MDS) classes form spatial niches. **A.** Schema and results of label transfer for cell classification of high-resolution spatial transcriptomic (stRNAish) datasets. **B.** Comparative coverage plot for the NF2 locus comparing Schwann^mer-^ and Schwann^mer+^ cells. **C.** Bar plot summarizing the NF2 gene expression across Schwann MDS classes in single cell and single nucleus RNAseq (sc/snRNAseq) datasets. **D.** Global chromatin accessibility (fragments) plotted in a standard Circos plot with segments representing chromosomes. Inner circle represents the excess accessibility found in Schwann^mer-^ cells. **E.** Geneset enrichment (GSEA) plot for pathways represented by RNA molecules enriched in Schwann nuclei (when compared with cytoplasmic RNA). **F.** Spatial Cellcharter cluster stability analysis for optimal spatial niches in stRNAish. **G.** Results of spatial niche classification of the stRNAish datasets.

**Sup. Fig. S6:**
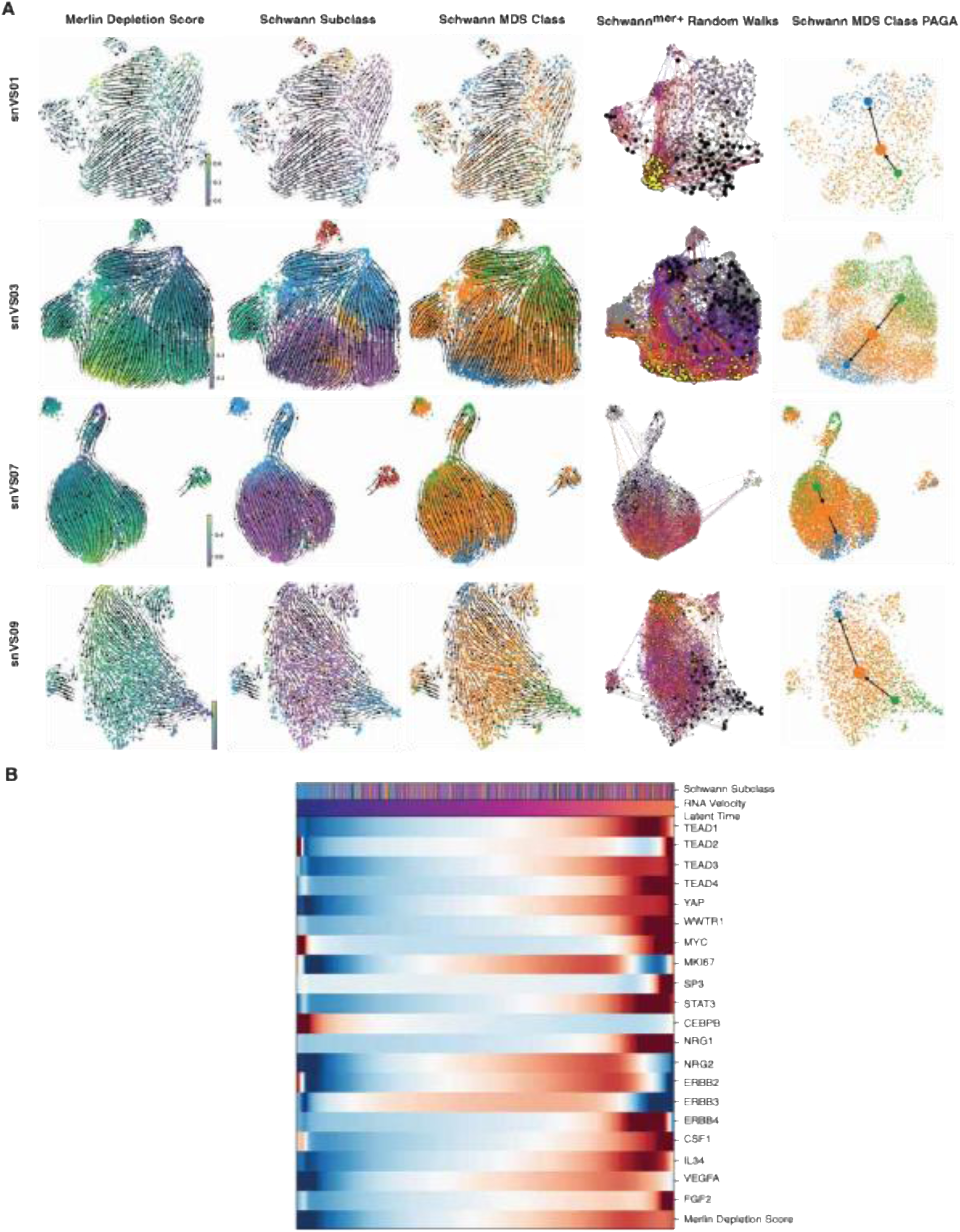
Schwann cell state plasticity in individual tumors. **A.** Left: Representative uniform manifold approximation and projection (UMAP) of Schwann cells from single tumor samples (VS01, VS03, VS07, VS09), overlaid with RNA velocity trajectories on merlin depletion score (MDS), Schwann sub-classes, and Schwann MDS class. Right: UMAP showing random walk trajectory based on RNA velocity of Schwann^mer+,^ with yellow dot representing Schwann^mer+^ cell’s inferred final state after 100 random walks. Far Right: Directional PAGA graph abstraction of RNA velocity across Schwann cells stratified by MDS status. **B.** A heatmap for tumorigenic genes and merlin depletion score arranged along the RNA Velocity Latent Time in Schwann cells.

**Sup. Fig. S7:**
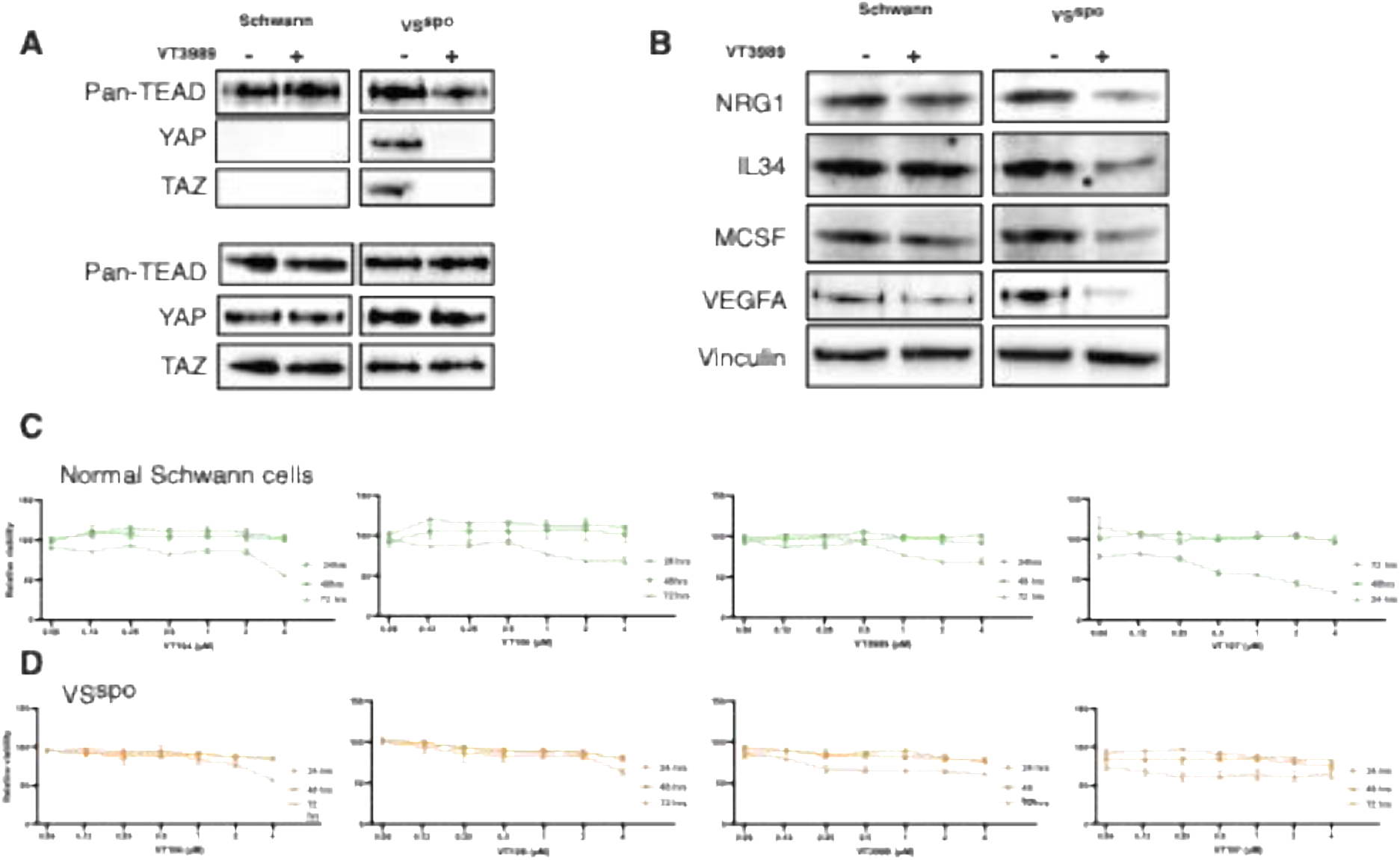
Pharmacological TEAD1 inhibition in human-derived tumor cells. **A.** Co-immunoprecipitation of human normal Schwann cells, and primary cells cultures from sporadic vestibular schwannoma (VS^spo^). The top section demonstrates the results from pan-TEAD immunoprecipitation with and without treatment with TEADi (VT3989; 4 µM, 6 hours) for each cell type. The bottom section demonstrates the blots from cell lysates (input), with GAPDH as loading control. **B.** Representative western blot of the 4 cell types as in E., treated with VT3989 (4 µM, 6 hours), and the effects on tumorigenic mediators (MCSF, VEGF-A, IL-34, and NRG1). Vinculin was used as loading control. **C.** and **D.** Comparative survival (CellTiter-Glo) of normal human Schwann cells (green), and schwannoma cells (Spo Swn; orange) in primary culture, with exposure to 4 TEADis (VT104, VT106, VT3989, and VT107), at 24, 48 and 72 hours, across concentrations (0.06 – 4 µM). Data presented as mean ± SEM.

